# Neural bases of reading fluency: A systematic review and meta-analysis

**DOI:** 10.1101/2023.09.25.559403

**Authors:** Marissa M. Lee, Catherine J. Stoodley

## Abstract

Reading fluency, the ability to read quickly and accurately, is a critical marker of successful reading and is notoriously difficult to improve in reading disabled populations. Despite its importance to functional literacy, fluency is a relatively under-studied aspect of reading, and the neural correlates of reading fluency are not well understood. Here, we review the literature of the neural correlates of reading fluency and rapid automatized naming (RAN), a task that is robustly related to reading speed. In a qualitative review of the neuroimaging literature, we evaluated structural and functional MRI studies of reading fluency in readers ranging in skill levels. This was followed by a quantitative activation likelihood estimate (ALE) meta-analysis of fMRI studies of reading fluency and RAN measures. We anticipated that reading speed, relative to untimed reading and reading-related tasks, would harness ventral reading pathways that are thought to enable the fast, visual recognition of words. The qualitative review showed that speeded reading taps the entire canonical reading network. The meta-analysis, which focused on in-scanner reading fluency measures, indicated a stronger role of the ventral reading pathway in fluent reading. Both reviews identified regions outside the canonical reading network that contribute to reading fluency, such as the bilateral insula and superior parietal lobule. We suggest that fluent reading engages both domain-specific reading pathways as well as domain-general regions that support overall task performance and discuss future avenues of research to expand our understanding of the neural bases of fluent reading.

## 1. Introduction

Reading is a complex task that requires processing of written language at multiple levels, from single letters to single words, to the understanding of syntax and ultimately comprehension. Behaviorally, reading and reading-related abilities have been studied from developmental time points from pre-readers to skilled readers, with heavy emphasis on phonological processing in typical reading development and reading disorders. Reading mastery is not defined solely through the fundamental process of decoding written text, but also incorporates the speed at which one can comfortably read and comprehend text. Throughout the reading literature, this speed-related skill has been referred to as reading speed, reading fluency, and even reading automaticity, which captures the fact that typical text decoding is often performed without conscious effort.

Reading fluency has been studied across many languages, ranging from shallow orthographies with easy to predict sound-to-letter mappings (e.g., German), to deep orthographies where the sound-to-letter mappings are harder to predict and include many irregularities (e.g., English). Measuring fluency in shallow orthographies is essential when determining reading disability, as measures of phonological skills often result in a ceiling effect due to the predictable relationship between letters and sounds (Ellis et al., 2004; Frith et al., 1998; Serrano & Defior, 2008; Wimmer, 1993; Wimmer & Mayringer, 2002). Deep orthographies tend to use a combination of phonological decoding and fluency measures to detect reading difficulties, but it is noteworthy that reading speed is usually impaired regardless of the orthographic depth of the language (Diamanti et al., 2017; Jednoróg et al., 2015; Landerl et al., 2013).

While clearly an important component of effective reading, the mechanisms and neural bases of reading fluency are relatively under-investigated compared with decoding skills. It is not clear whether fluent reading requires only an optimally-functioning reading network or whether reading that is both accurate and fast taps additional neural resources that can be harnessed to improve reading speed. Here, we evaluate the current state of our understanding of the neural bases of reading fluency with both a qualitative review and a quantitative meta-analysis of functional imaging studies of reading speed and fluency-related tasks.

## 2. The Reading Network

Reading incorporates a network of cortical and subcortical regions, harnessing areas involved in visual processing, language processing, and the integration of the two. Three cortical areas consistently emerge as hubs of the reading network: the left inferior frontal gyrus (IFG), left occipito-temporal (OT) cortex, and left temporo-parietal cortex (TP; sometimes referred to as the temporoparietal junction [TPJ]; **Figure 1**; Kearns et al., 2019; Linkersdörfer et al., 2012; Maisog et al., 2008; Martin et al., 2015; Pollack et al., 2015; Richlan et al., 2009). Each of these regions has been associated with different aspects of reading and tend to show differential engagement in cohorts with diagnosed reading disability (Martin et al., 2016; Richlan et al., 2009; Richlan et al., 2012).

**Figure 1.**
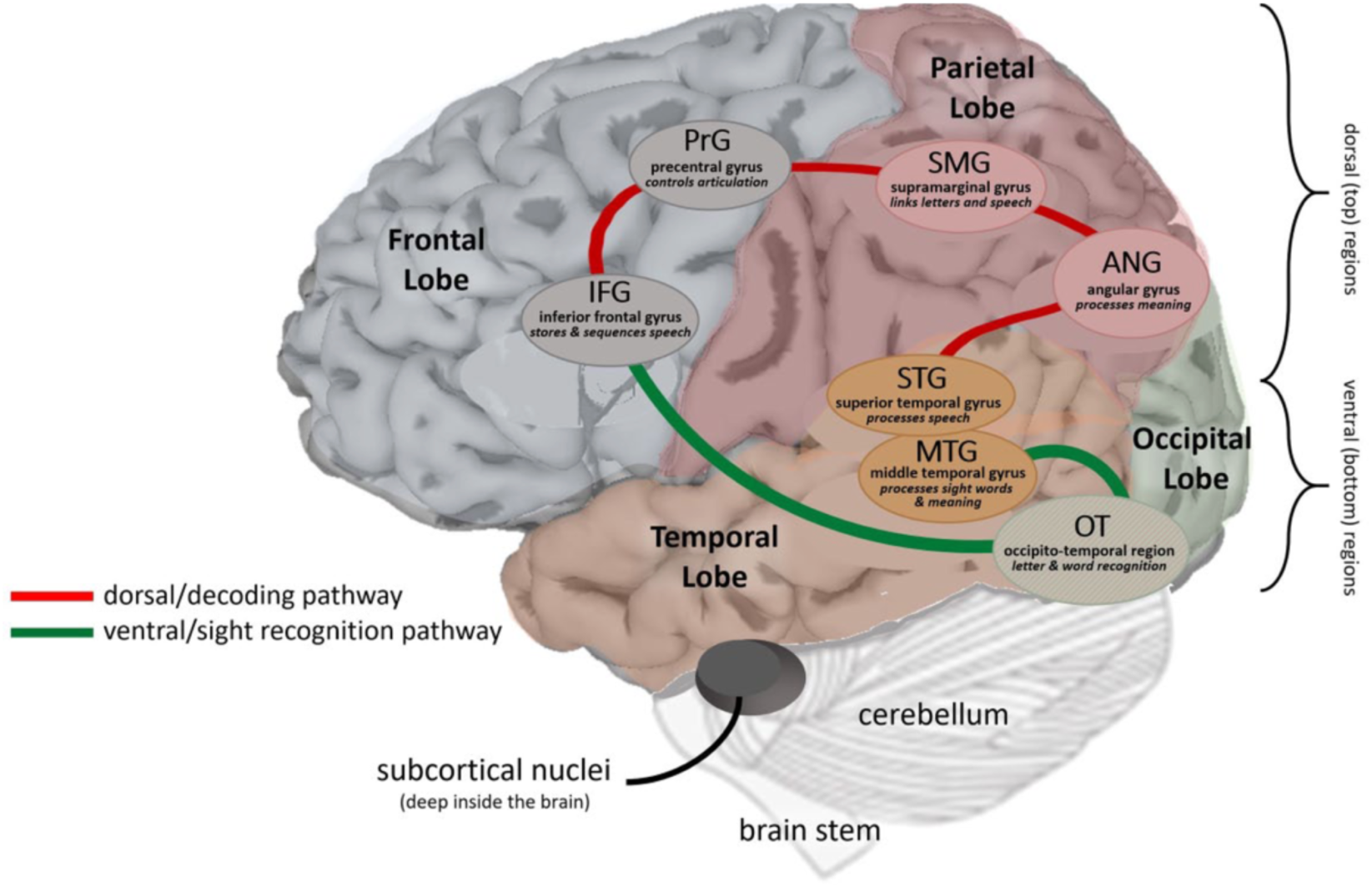
The canonical reading network (with permission from Kearns et al., 2019)

The reading-related regions of the left IFG overlap with Broca’s area (Brodmann areas [BA] 44, 45, 47), a region historically connected with language production. More recent research shows the left IFG to be involved in phonological, syntactic, and semantic language processes. Left IFG activation appears early in word processing, which may reflect early phonological processing (Cornelissen et al., 2009). A study of pre-readers also suggested that left IFG supports phonological processing and naming speed via functional connections with other cortical reading network regions (Benischek et al., 2020). There is also evidence of left IFG activation when evaluating the semantic (Acheson & Hagoort, 2013; Chou et al., 2009) and syntactic information (Acheson & Hagoort, 2013) of a word. There is an anterior-to-posterior functional gradient in the IFG, such that the anterior IFG (BA 47) is involved in sentence-level and semantic processing, syntactic processing engages the junction of BA 44/45, and word-level and phonological processing is represented more posteriorly (BA 44; Uddén & Bahlmann, 2012).

The left OT includes the lingual gyrus and fusiform gyrus, which are associated with word recognition. Activation of the left lingual gyrus, also called the medial OT gyrus, has been associated with the visual recognition of word length (Mechelli et al., 2000). It has been proposed that this pattern reflects the role of the lingual gyrus in processing global aspects of visual information when reading rather than local information (Fink et al., 1996), though it should be noted that the processing of local information was associated with engagement of the right lingual gyrus. The ventral OT (vOT) cortex, a region of the left fusiform gyrus that is often called the visual word form area (VWFA) for its sensitivity to word recognition (Wandell et al., 2012; Weiner & Zilles, 2016), shows higher engagement when viewing words compared to false fonts or line stimuli, even when viewing words passively rather than completing a reading or language task (Ben-Shachar et al., 2007). Therefore, this region shows specialization for words compared to other visual stimuli in literate individuals.

The left TP areas involved in reading include the supramarginal gyrus (SMG), angular gyrus (AG), and posterior regions of the middle and superior temporal gyri (MTG/STG). The primary roles of the left SMG during reading seem to be orthographic (González-Garrido et al., 2017; Xia et al., 2018) and phonological processing (Dickens et al., 2019; Sliwinska et al., 2012; Xia et al., 2018). The left AG has been associated with phonological processing (Benischek et al., 2020) and semantics (Paz-Alonso et al., 2018; Seghier, 2013). Lastly, the STG overlaps with Wernicke’s area, another traditional language region of the brain, which is involved in language comprehension. This region is also associated with reading-related processes such as phonological awareness (Wang et al., 2020; Xia et al., 2018), semantics and syntax (Vigneau et al., 2006), and audiovisual integration during speech processing (Ye et al., 2017).

Subcortically, the thalamus is consistently activated during reading tasks, but is not usually included as part of the reading network. The structure and function of the left thalamus has been associated with single word reading (Benischek et al., 2020; Lebel et al., 2013; Lee et al., 2023) and reading fluency (Lebel et al., 2013), and there are structural and functional differences in the thalamus between individuals with typical reading development and those diagnosed with Developmental Dyslexia (DD, also known as reading disorder [RD] or Specific Learning Disorder with Impairment in Reading; Diaz et al., 2012; Jednoróg et al., 2015; Maisog et al., 2008; Richlan et al., 2009).

Lastly, the cerebellum is also consistently engaged during reading tasks, but, like the thalamus, is rarely included as part of the canonical reading network. Right crus I of the cerebellum interconnects with left hemisphere regions supporting reading and is activated in concert with these regions during reading tasks, as evident from a neuroimaging meta-analysis of typical adult readers (Martin et al., 2015). Functional and structural differences between readers with typical development and those diagnosed with DD have also been found in the posterolateral cerebellum (e.g., crus I, crus II, lobule VI; Greeley et al., 2020; Stoodley, 2014; Stoodley & Stein, 2011).

The aforementioned regions comprising the cortical reading network can also be divided into the dorsal (**Figure 1, red**) and ventral reading pathways (**Figure 1, green**; Kearns et al., 2019). TP regions (left STG, AG, SMG) and the IFG make up the dorsal reading pathway, which generally supports decoding words. In typical reading development, this often corresponds to sounding out individual letters (sound-to-letter mapping) in order to decode the whole word. The dorsal reading pathway overlaps with the dorsal *visual* pathway, which has been the target of transcranial magnetic stimulation (TMS) research showing a relationship between posterior cortical regions and eye movements associated with reading (e.g., saccades; Laycock & Crewther, 2008). The ventral reading pathway includes OT regions such as the VWFA, the MTG, and the IFG and is associated with rapid recognition of whole words, usually already-familiar sight words that do not need to be decoded. The ventral pathway is used for quick and efficient reading, as visually identifying a whole word takes less time than linking individual letters to phonemes to sound out a word.

Anatomical connectivity of white matter (WM) pathways has been associated with reading ability in diffusion tensor imaging (DTI) studies. Multiple WM tracts correlate with performance on reading tasks, and, similar to gray matter (GM) findings, these tracts tend to be left-lateralized and are consistent with the subdivision of the dorsal and ventral reading pathways. WM tracts can be divided into pathways supporting phonological processing or those involved in the orthographic (or visual) processing of words. The main tract associated with phonological processing is the arcuate fasciculus (Dick & Tremblay, 2012; Yeatman et al., 2012), considered part of the superior longitudinal fasciculus (SLF; Dick & Tremblay, 2012), which connects Broca’s and Wernicke’s areas. The middle longitudinal fasciculus (MLF) has also been associated with phonological processing and connects the AG with the STG (Dick & Tremblay, 2012). The main WM tract supporting orthographic processing is the left inferior longitudinal fasciculus (ILF), which connects occipital regions to superior temporal cortical regions (Dick & Tremblay, 2012; Yeatman et al., 2012), including the vOT, which is associated with rapid word recognition and includes the VWFA.

The left internal capsule (Beaulieu et al., 2005), left centrum semiovale (CS) and the left superior corona radiata (SCR; Niogi & McCandliss, 2006) have also been associated with reading ability, specifically with untimed word identification tasks. The fibers composing each of these tracts flow into each other, with the centrum semiovale being the most lateral, the internal capsule being the most medial and surrounding the basal ganglia, and the corona radiata located in between. The anterior internal capsule carries information from the thalamus through the corona radiata and centrum semiovale to the prefrontal cortex while the posterior internal capsule carries information from the thalamus to the cerebellum (Emos & Agarwal, 2021). Both pathways may be important in relaying reading information between these brain regions that have been implicated in single word reading and reading fluency. The anterior internal capsule also carries information from the caudate (Emos & Agarwal, 2021) which, as will be discussed below, has been associated with reading fluency.

A review of WM tracts supporting reading and language (Dick & Tremblay, 2012) additionally identified the uncinate fasciculus, which connects anterior temporal regions to inferior frontal regions, and the extreme capsule, which connects inferior frontal regions to middle and superior temporal cortices, as being associated with semantic processing. The extreme capsule, along with the MLF, has also been associated with the ventral language pathway which supports “sound-to-meaning” processing.

In summary, the reading network consists of largely left-lateralized regions involved in the visual processing of letters (e.g., vOT), phonological processing (e.g., IFG), understanding the syntax of full sentences, and the semantic meaning of words (e.g., TP regions). Evidence suggests that subcortical structures support the cortical reading network, though specific contributions of regions like the thalamus and cerebellum are still unclear.

## 3. Behavioral measures of reading fluency

Multiple tasks have been used to measure the fluency or automaticity of reading. The Test of Word Reading Efficiency (currently in its second edition; TOWRE-2; Torgesen et al., 2012) measures speed and accuracy of reading out loud under a time constraint. The TOWRE-2 is composed of two sub-tasks testing speeded reading of single real words (Sight Word Efficiency; SWE) and nonwords (Phonemic Decoding Efficiency; PDE). Both SWE and PDE consist of lists of words or nonwords that are read out loud as quickly and accurately as possible. The words increase in difficulty as the lists progress. Individuals are scored based on how many words or nonwords are read accurately within 45 seconds. SWE was designed to test an individual’s ability to recognize whole words fluently and automatically by sight as opposed to focusing on single letters; and PDE targets the ability to blend phonemes together, which is necessary for fluent reading.

Reading speed can also be measured by having participants quickly read sentences and passages followed by questions to assess comprehension. For example, in the Gray Oral Reading Test (GORT-5; Wiederholt & Bryant, 2012), the amount of time it takes to read a short story and the accuracy of reading are combined to create a fluency score. Research groups have also created their own sentences or passages to administer to participants with instructions to read as quickly as possible. These stimuli are also usually followed by questions assessing comprehension (e.g., about content or if a sentence made grammatical sense) to confirm that increases in speed did not decrease reading accuracy. Another option some research groups have adopted is the manipulation of word or sentence presentation rates. With this method, the speed of sentence and passage reading is controlled by displaying one word of a sentence at a time to establish a comfortable reading rate specific to each participant. This speed can then be modified to create slower (“constrained”) and faster (“accelerated”) rates of reading (e.g., Langer et al., 2015).

The most common non-reading task that is associated with reading fluency is rapid automatized naming (RAN), in which participants name a list of highly familiar items as quickly as possible. RAN tasks can be composed of colors, objects, digits, or letters, and can be administered to participants of a wide range of ages and abilities. RAN is included as a subtest in multiple literacy assessment batteries, including the Woodcock Reading Mastery Tests (WRMT-III; Woodcock, 2011) and the Comprehensive Test of Phonological Processing (CTOPP-2; Wagner et al., 2013). The direct relationship between rapid naming and reading is not entirely understood, but it has been suggested that rapid naming represents a “microcosm” of the overall reading network (Norton & Wolf, 2012) and RAN performance predicts reading fluency across many orthographies (Georgiou et al., 2007). Importantly, the ability of non-alphanumeric RAN (i.e., objects, colors) to predict reading scores tends to decrease as children age, as these stimuli tend to take more time to name than the increasingly familiar alphanumeric stimuli. The correlation between alphanumeric RAN and reading ability persists throughout the lifespan (Norton & Wolf, 2012). RAN may additionally be a predictor of general processing speed ability, which is also related to reading fluency and may be a contributing factor to poor reading fluency skills in those diagnosed with DD, though this is still debated (Norton & Wolf, 2012; Wolf & Bowers, 1999). Research has shown a relationship between faster RAN scores and faster processing speed ability (Gerst et al., 2021), and slower processing speed ability and reading fluency has been identified in clinical groups with no diagnosed reading difficulties (Jacobson et al., 2011; Rucklidge & Tannock, 2002). Behaviorally, though, the evidence connecting general processing speed ability with reading speed has been inconsistent (Norton & Wolf, 2012).

## 4. The neural bases of reading fluency

There is an extensive neuroimaging literature investigating the neural correlates of reading in individuals with typical reading development as well as those with DD, but most studies focus on phonological and single-word decoding skills. A previous review emphasized the importance of studying reading fluency and provided a deep dive into the literature surrounding the use of RAN (Norton & Wolf, 2012). Here, we focus on recently published neuroimaging work examining reading fluency. It is important to note that “reading fluency” has many definitions; our use of the term “reading fluency” or just “fluency” will focus on the speed of accurate reading.

### 4.1 Structural MRI

#### 4.1.1 Gray matter studies

A handful of studies have used structural MRI techniques to investigate the GM correlates of reading fluency in both typical readers (He et al., 2013; Houston et al., 2014) and reading impaired populations (Fernandez et al., 2013; Jednoróg et al., 2015; Kronbichler et al., 2008; Liu et al., 2013; Steinbrink et al., 2008). Houston and colleagues (2014) evaluated the relationship between GM volume and reading fluency and rapid naming at two timepoints in adolescent English readers: first at baseline and again approximately two years later. They found that higher reading fluency scores at baseline predicted decreases in GM volume between baseline and follow-up in the left inferior parietal cortex, and better rapid naming scores at baseline predicted decreases in GM volume between baseline and follow-up in the left inferior parietal and inferior frontal gyrus. Contrasting results were found in adult native Chinese speakers with English language education (He et al., 2013), where positive correlations were reported between GM volume and both Chinese and English reading fluency measures in multiple brain regions commonly associated with the reading network (e.g., left AG, STG), but no negative correlations were found. In addition, GM volumes of the left posterior precuneus, right insula, right middle frontal gyrus (MFG), and left anterior cingulate cortex (ACC) showed high predictive accuracy for fluency scores. These differences could be due the age of the participants—adolescents are still undergoing developmental changes in GM, especially in inferior parietal and frontal brain regions where GM has peaked and started to decrease, whereas adults show more stable GM trajectories (Giedd & Rapoport, 2010).

There are inconsistent findings when comparing GM correlates of fluency in individuals with typical reading development compared with those diagnosed with DD, though most studies converge to show structural correlates of fluency in the temporal cortex, TPJ, and OT regions. In Germans with typical reading development and DD readers, Kronbichler and colleagues (2008) found positive correlations between the number of sentences read in one minute and GM volume in regions where impaired readers had significantly less GM compared to controls, namely in bilateral OT cortices and the right SMG. A study investigating reading across languages (German, French, and Polish) found, regardless of language or reading ability, a significant positive correlation between right superior temporal pole GM volume and RAN-pictures performance, and negative correlations between GM volume and RAN-pictures in left lingual gyrus, left postcentral gyrus, right precentral gyrus, right MFG, and left insula. Researchers also found correlations within the right precentral gyrus that differed between typical and DD cohorts. Typical readers showed a negative correlation between right precentral gyrus volume and RAN-pictures, whereas DD readers showed a positive correlation (Jednoróg et al., 2015). Conversely, Steinbrink and colleagues (2008) found pseudoword reading speed correlated with GM volume in the left STG only in German readers with typical reading acquisition; the same pattern was not evident in the DD cohort. GM in the left anterior STG has also been associated with character reading fluency in Chinese DD readers (Liu et al., 2013).

Studies have also reported cerebellar correlates of reading fluency measures. Along with OT and temporal cortical regions, Kronbichler and colleagues (2008) found positive correlations between the number of sentences read in one minute and GM volume in bilateral cerebellar regions (lobule VI). Jednoróg and colleagues (2015) also reported that right lobule VI GM correlated with RAN-pictures scores. Fernandez and colleagues (2013) focused solely on the cerebellum’s involvement in reading and divided their DD group into those with only impaired decoding and those with only impaired fluency. In the impaired decoding group, they found positive correlations between fluency measures and the left inferior-posterior cerebellar region of interest (ROI) which included Crus II, lobule VIIB, and lobules VIII-X. The impaired fluency group showed positive correlations between scores on a single word reading task and total cerebellar GM volume and total overall cerebellar volume.

GM regions associated with fluency scores in structural MRI studies overlap heavily with the traditional reading network and include the cerebellum. Additional regions outside the reading network that have been associated with fluency measures include the precuneus, cingulate cortex, and insula. These regions are mostly left-lateralized and may either support reading or assist in directing attention during reading tasks. In summary, increases in GM tend to correlate with better reading fluency in non-DD readers, while the correlations in DD readers sometimes deviate from this relationship. Age is also an important factor when considering the direction of the correlation between GM and fluency measures.

#### 4.1.2 White matter studies

White matter (WM) correlates of reading fluency have also been investigated in typically developing readers (Lebel et al., 2013, 2019; Rimrodt et al., 2010; Rollans et al., 2017; Steinbrink et al., 2008; Tschentscher et al., 2019) and DD (Lebel et al., 2019; Rimrodt et al., 2010; Steinbrink et al., 2008; Tschentscher et al., 2019). Regardless of reading ability, fluency has been positively correlated with WM fractional anisotropy (FA) in the bilateral IFG, left lingual gyrus, left fusiform gyrus, and left MTG in youths, which overlaps extensively with the typical reading network (Rimrodt et al., 2010). In groups with typical reading development, converging findings indicate that WM in the SLF, ILF, and corona radiata correlate with fluency, as discussed further below. All of these pathways have been previously associated with reading and language skills (Dick & Tremblay, 2012; Takeuchi et al., 2016; Yeatman et al., 2012). The correlation between fluency scores and the SLF likely reflects the tract’s role in linking the frontal, temporal, and parietal lobes, all of which are involved in the reading network, while the ILF’s connections between temporal and occipital lobes are consistent with the reported role of the OT in reading fluency and sight word recognition. Increases in FA in both the SLF and ILF has been associated with literacy acquisition in younger individuals, while the ILF has also been correlated with maintaining reading skills later in life (Cheema et al., 2018). The link between the corona radiata and reading possibly stems from its role in linking the thalamus and cortex; thalamic GM is also associated with reading scores (e.g., Lee et al., 2023).

Lebel and colleagues (2013, 2019) have reported positive relationships between reading fluency and FA of the bilateral SLF, bilateral inferior longitudinal/fronto-occipital fasciculus, bilateral posterior corona radiata, and right anterior corona radiata, along with bilateral regions of the corpus callosum and right anterior limb of the internal capsule. Focusing on the different WM correlates of various RAN tasks, Rollans and colleagues (2017) also reported correlations between RAN-objects scores and the left ILF and SLF; between RAN-digit scores and the right superior corona radiata; and between RAN-letters scores and the right anterior corona radiata. Tschentscher and colleagues (2019) found positive associations between RAN-letter and RAN-digit naming speed and structural connectivity between the planum temporale and medial geniculate body of the thalamus in typically developing readers, but this relationship was not seen in a DD cohort. Whole-brain DTI studies correlating fluency scores with WM reveal negative relationships with FA in the right frontal lobe (Rollans et al., 2017; Steinbrink et al., 2008) including the IFG and prefrontal cortex (PFC), as well as the right temporal lobe and left occipital lobe (Rollans et al., 2017).

Lebel and colleagues (2019) investigated the trajectory of WM development in DD youths who were either impaired in both fluency and decoding or impaired in only fluency by collecting DTI data on two occasions approximately two years apart. Youths impaired in both fluency and decoding showed a decrease of mean diffusivity (MD) of the right anterior corona radiata between baseline and the second visit, whereas both groups showed MD decreases in the left uncinate fasciculus and right posterior corona radiata. Steinbrink and colleagues (2008) found negative relationships in adults with a childhood diagnosis of DD between reading speed performance on a pseudoword reading task and FA in the left external capsule, left STG, and left middle occipital gyrus (MOG), overlapping with the left SLF.

While the WM regions and tracts identified in these studies correspond with pathways that are part of the reading network, there are still questions that remain. WM tract findings are not as left-lateralized as cortical GM findings are, suggesting that the right hemisphere may play a larger role in reading fluency than the GM results indicate. Also, while most studies report increased structural connectivity associated with better fluency scores, there are also reports of reduced structural connectivity or reduced FA associated with better fluency, which seems counterintuitive. Reading ability and developmental stage may be a factor in these discrepancies.

### 4.2 Functional MRI

Functional MRI studies have contributed greatly to our understanding of the neural correlates of reading fluency, with studies investigating many types of fluency tasks in languages with both shallow (or transparent) and deep (or opaque) orthographies. Converging evidence supports strong engagement of the left-lateralized reading network, including inferior frontal, TP, and OT regions during reading fluency tasks, with trends toward greater activation of the network as reading fluency increases.

#### 4.2.1 Task-based (fMRI) studies

In youths and young adults, performance on rapid naming tasks has been positively associated with activation in left IFG, left inferior temporal gyrus (ITG), bilateral MFG, right STG, right superior temporal sulcus (STS), right MTG, left SMG, left AG, and right cerebellum (Misra et al., 2004; Turkeltaub et al., 2003), aligning with the established reading network. Tasks that control the speed of reading (e.g., comfortable pace, slowly, and rapidly) result in a gradient of activation in the left fusiform gyrus and the surrounding OT region, where accelerated reading shows the most activation and slow reading the least. Greater activation of the left fusiform gyrus was also apparent in fluent word reading compared to rapid letter naming, along with the left IFG and MTG (Benjamin & Gaab, 2012).

The left vOT / VWFA has been used as a region of interest (ROI) in multiple fMRI studies of fluency. The VWFA has been associated with word recognition (Dehaene & Cohen, 2011) which is essential for rapid reading. Letter fluency scores in English-speaking children, measured with rapid letter naming, negatively correlated with activity in the anterior vOT during an auditory phonological task but not with posterior vOT, a sub-region mostly recruited in adults (Wang et al., 2021). An effect of reading speed on vOT activation has also been found in French children first learning how to read: faster reading scores were associated with increased vOT activation when viewing everyday objects, faces, and words (Dehaene-Lambertz et al., 2018). This effect was strongest for the word reading condition, as opposed to viewing houses or faces, suggesting that vOT plays a specialized role in the rapid recognition of words. These two studies resulted in opposing directions of activation within the vOT, which could be due to the type of task completed in the scanner (auditory vs. visual). It is possible that vOT activation is higher at early stages of learning how to read when a word is visually presented versus auditorily presented.

Task-based fMRI research has also parsed brain activation patterns during different types of rapid naming tasks. Two studies compared RAN-letters, RAN-digits, RAN-objects, rapid word reading, and rapid pseudoword reading, showing both overlapping regions and brain activation patterns that vary depending on stimulus type (Cummine et al., 2014, 2015). For example, all alphanumeric (letter and digit) stimuli engaged the bilateral precentral gyrus, left middle occipital gyrus, right lingual gyrus, right superior frontal gyrus (SFG), right cerebellar lobule VI, and left precuneus (Cummine et al., 2015), whereas greater activation in alphanumeric compared to non-alphanumeric stimuli was found in traditional reading network areas (e.g., left STG, right lingual gyrus) and subcortical regions such as the bilateral thalamus, bilateral caudate, and left globus pallidus (Cummine et al., 2014). Rapidly reading real words resulted in increased activation within the right lingual gyrus, left MTG, and left cerebellar lobule VI compared to rapidly naming single letters, whereas reading pseudowords resulted in decreased activation in the right thalamus compared to single letters (Cummine et al., 2015). These findings suggest that all rapid naming and reading tasks recruit regions involved in articulation (e.g., precentral gyrus; Park et al., 2018) even when reading silently, but alphanumeric characters and words recruit more typical reading network regions. Consistent with this pattern, rapidly reading real words resulted in increased activation in the left MTG compared to identifying single letters. The left MTG has been associated with processing syntactic information during reading (e.g., Rodd et al., 2010; Ye et al., 2010), which is impossible to extract from single letters.

Relative to typical readers, DD readers tend to show decreased activation in brain regions engaged during fluent reading. When controlling sentence reading speed (slow, comfortable, or accelerated rate), DD readers showed a gradient effect in left SFG, right MFG, bilateral insula, bilateral MTG, left postcentral gyrus, left superior parietal lobule (SPL), bilateral inferior parietal lobule (IPL), right precuneus and left thalamus, where accelerated reading resulted in the highest amount of activity (Christodoulou et al., 2014). Regardless of reading rate, DD readers consistently showed reduced activation in these brain regions compared to those with typical reading development. In a similar task where sentences were read at either comfortable or accelerated rates (Langer et al., 2015), there was reduced activation in the DD group in the canonical reading network regardless of reading rate (left IFG, bilateral IPL, and left SMG; Langer et al., 2015). Similar results are also evident in native German speakers, where reading disability is defined primarily by poor fluency rather than phonological impairments. Dysfluent readers showed reduced activation in the left posterior MTG and left SMG, but also showed increases in activation in bilateral lingual gyrus, left medial temporal regions, left IFG, bilateral thalamus, and right caudate (Kronbichler et al., 2006). Furthermore, activity in the left SMG predicted reading fluency in DD, where higher rates of activity correlated with better fluency (Ozernov-Palchik et al., 2021). Brem and colleagues (Brem et al., 2020) also found a positive relationship between reading fluency and activity in the bilateral fusiform gyrus in impaired readers, again showing a connection between increased brain activity and increased fluency. Direct comparisons between typical and DD groups consistently report decreased activation in these regions in the DD group. Finally, right cerebellar lobule VI was the only brain region where activation patterns were associated with fluency ability: typical readers showed greater activation than children with impaired fluency (as indicated by RAN scores), and children with lower performance on *both* RAN and phonological tasks showed less engagement of right cerebellar VI compared to the impaired fluency-only group (Norton et al., 2014).

Taken together, DD groups tend to show reduced activation in left-lateralized cortical reading-related regions during fluency tasks compared to their typically reading counterparts, even when groups are matched for various demographic and non-reading cognitive abilities such as memory and general intelligence. There also seems to be a trend of increased use of right hemisphere reading network homologs in dysfluent readers compared to typical readers, suggesting the dysfluent readers may recruit these regions to compensate for the decreased engagement in left-hemispheric regions. These patterns also appear in pre-readers (Benischek et al., 2020), signifying that decreased activation and reduced connectivity within the reading network and recruitment of regions outside the reading network in those with reduced pre-reading rapid naming (e.g., RAN-colors) may be associated with later reading fluency outcomes.

#### 4.2.2 Functional connectivity (fcMRI) studies

Resting-state fMRI (rs-fMRI) research has not focused on reading fluency as much as task-based fMRI, but some insights can be gleaned from studies conducted primarily in shallow orthographies that rely more heavily on fluency rather than phonology to categorize DD. A whole-brain analysis of native Spanish speakers used independent components analysis (ICA) to identify a reading-related network. Functional connectivity (FC) within the reading-related ICA component positively correlated with reading fluency scores, specifically FC between left frontal regions, STG, MTG, thalamus, and caudate (Alcauter et al., 2017). A study conducted in native Chinese speakers reported similar findings, revealing that the connectivity strength between left MFG and the intraparietal sulcus (IPS) positively correlated with reading fluency (Zhou et al., 2016).

Reduced FC can also be seen in dysfluent readers when directly compared to those with typical reading development, especially in the reading network. When native Japanese speakers read in Hiragana, a Japanese syllabary with a very shallow orthography, individuals with DD (characterized by poor fluency) showed decreased FC between a seed region in the left fusiform gyrus and left ventrolateral prefrontal cortex (VLPFC), TPJ, IPL, MTG, insula, medial PFC, and right temporal pole, but increased connectivity with the left precentral gyrus, compared to typical readers (Hashimoto et al., 2020). A comparison of high- and low-fluency readers in English also showed reduced FC in low-fluency readers within two cognitive control networks: the cingulo-opercular (CO) and ventral attention (VA) networks (Freedman et al., 2020), the latter of which overlaps extensively with the traditional reading network’s frontal and temporal regions. Last, evidence of reduced FC between the cerebellum and cortical reading regions has been reported in individuals diagnosed with DD (Greeley et al., 2020). Reading fluency scores in the DD group positively correlated with FC between right cerebellar lobule VIII and the left AG. Further, reading fluency scores predicted the strength of FC between a cerebellar network identified through ICA and the right precentral gyrus (Greeley et al., 2020).

Across multiple languages, FC patterns associated with reading fluency overlap extensively with the traditional reading network, along with subcortical regions such as the cerebellum and thalamus. Dysfluent readers tend to show decreased FC between reading network regions, aligning with task-based results. This suggests that, along with decreased activation in reading-related brain regions, there is also less efficient communication between these regions in dysfluent readers.

### 4.3 Intervention and neuromodulation to increase reading fluency

Fluency training has been implemented in the hopes of increasing reading speed while not sacrificing accuracy, and neuroimaging has been used to understand how these interventions impact the neural underpinnings of reading. Neuromodulation studies have also investigated how directly modulating the brain impacts reading fluency, providing further insight into the neural correlates of fluent reading.

#### 4.3.1 Readers with typical reading development

To date, reading speed training for typical readers has been investigated at the neural level in English (Ferguson et al., 2014) and Japanese (Fujimaki et al., 2004, 2009). In English, participants were invited to complete a 6-week intervention where training focused on incrementally increasing reading speed by presenting passages at slow, medium, and fast rates, and training visual attention to a geometric shape placed at different locations on a computer screen at varying rates (Ferguson et al., 2014). Behaviorally, the training program increased reading speed. Ferguson and colleagues (2014) then chose bilateral inferior frontal and superior temporal regions (Broca’s and Wernicke’s areas in the left hemisphere) as regions of interest to determine if any resting-state changes occurred in these reading-related areas. While a decrease in FC between Broca’s area and its right hemisphere counterpart was seen after training was complete, this change in FC did not correlate with changes in reading speed rates (Ferguson et al., 2014).

Fujimaki and colleagues (2004, 2009) trained native Japanese speakers in the Park-Sasaki method (described in Miyata et al., 2012) which involves a combination of visual training and maintaining a meditative state. This can allow one to read up to 10,000 characters per minute when fully trained without affecting comprehension (Fujimaki et al., 2009). During neuroimaging, individuals trained or not trained in the Park-Sasaki method read a novel at either a comfortable pace or as rapidly as possible while still maintaining comprehension of the text. When reading rapidly, trained individuals tended to show decreased activation in Broca’s and Wernicke’s areas (Fujimaki et al., 2004, 2009), along with decreased activation in additional left-lateralized frontal and superior temporal regions. Interestingly, there was a mixture of increases and decreases in activation after training in the right hemispheric homologs of the reading network (Fujimaki et al., 2009). Overall, regardless of training, brain activation in left lateralized reading-related regions tended to decrease during rapid reading compared to reading at a comfortable pace.

Taken together, fMRI research suggests behavioral intervention targeting reading speed is associated with decreases in activation, especially in regions overlapping with Broca’s and Wernicke’s areas. One could argue that behavioral fluency training affects the reading network by requiring less engagement as the reading task becomes more automated. Of note, the above studies restricted data analyses to very specific regions within the reading network by choosing an ROI approach, so it is possible that other brain regions show relevant fluency-associated activation changes.

#### 4.3.2 Readers diagnosed with reading disability

Similar to research in typical readers, the neural underpinnings of fluency training in DD have not been studied extensively, but there are two studies in English (Horowitz-Kraus et al., 2014; Rezaie et al., 2011) and one in French (Stappen et al., 2020) which examined the neural correlates of fluency training in youth with reading difficulties.

Rezaie and colleagues (2011) focused on training native English-speaking DD readers in word study (i.e., learning patterns within words, as opposed to rote memorization), fluency, vocabulary, and comprehension over the course of one year. MEG data were acquired at baseline. After training, the impaired readers were split into either adequate responders or inadequate responders depending on their fluency improvements. The baseline MEG recordings indicated that post-training reading fluency gains were predicted by activity in left IPL (AG and SMG) and superior temporal gyrus (STG): increased activity in these brain regions correlated with bigger gains in fluency between baseline and post-training in poor readers. This suggests that brain activation patterns pre-intervention can predict later fluency performance, and, more specifically, that higher levels of activation within the left IPL and STG are predictors of greater improvement in fluency.

Neural changes after exposure to the Reading Acceleration Program (RAP) have also been studied in native English speakers (Horowitz-Kraus et al., 2014). RAP focuses on improving reading fluency through word decoding and comprehension, where task speed is increased as the participant improves. fMRI scans were conducted at baseline and after four weeks of RAP training to test the effects of RAP on reading and executive control network ROIs. Researchers found a positive correlation between an improvement in reading fluency scores and activation in left ACC, left MFG, left fusiform gyrus, and left inferior occipital gyrus, showing a relationship between improved fluency and increased engagement of both the executive control network and regions of the ventral reading network.

Last, the neural correlates of RAN training have been studied in native French-speaking children diagnosed with DD (Stappen et al., 2020). The RAN intervention focused on increasing the speed of naming colors and objects from a broad range of categories (e.g., home objects, animals). FA values in white matter tracts of interest were calculated at baseline and after RAN training was complete, approximately eight weeks later. Improvements in RAN scores positively correlated with increased FA in the left anterior arcuate fasciculus, suggesting strengthening of this pathway.

Interestingly, fluency interventions for DD readers more often resulted in increases of activity associated with gains in fluency, whereas typical readers show decreases in activation when reading more rapidly. This begs the question as to whether decreases in activity should be interpreted as atypical or as evidence of increased efficiency. It is possible that the neural mechanisms underpinning changes in reading fluency differ depending on the initial fluency levels of the participants.

### 4.4 Neuromodulation

#### 4.4.1 Transcranial direct current stimulation studies

Transcranial direct current stimulation (tDCS) is a neuromodulation method in which two electrodes apply a small amount of electrical current (1-2 mA) directly to the scalp for a short amount of time (approximately 20 minutes). TDCS utilizes an anode and cathode for three modes of operation: facilitating brain activity (anodal), inhibiting brain activity (cathodal), or a sham condition in which there are no neuromodulatory effects (Ferrucci et al., 2015; Woods et al., 2016). It has been suggested that those diagnosed with DD might benefit more from tDCS compared to those with typically developed reading skills, because the latter group has already attained proficient reading skills with no real room for improvement (Cancer & Antonietti, 2018).

The effects of tDCS in typical and DD groups have been mixed, in part because multiple different protocols have been used (e.g., varying in the amount of voltage, duration, reference electrode placement, and brain regions targeted). Turkeltaub and colleagues (Turkeltaub et al., 2012) utilized tDCS to facilitate neuronal activity in the left posterior temporal cortex (overlapping with MTG and STG) while inhibiting activation in the right hemisphere homologue to investigate the impact of left-lateralizing activity on reading ability. Modulating this region resulted in improved timed reading scores, indicating that increasing left lateralization of posterior temporal cortical activation can lead to more efficient reading. Interestingly, shifting electrodes superiorly to the TPJ showed the opposite effect: facilitating activation in the right hemisphere resulted in increased fluency compared to facilitation of the left hemisphere (Thomson et al., 2015). Cathodal tDCS applied to the cerebellum (Boehringer et al., 2013) and anodal tDCS applied to the left IFG and MTG (Westwood et al., 2017) did not affect reading fluency in typical readers.

Additional tDCS studies have been carried out in DD cohorts to explore neuromodulation as a potential intervention or treatment. The greatest success has come from targeting the left TPJ with anodal tDCS (for review see Cancer & Antonietti, 2018), though effects were different in adults compared to children and adolescents. Younger samples improved on nonword fluency tasks whereas adults improved in real word fluency. Lazzaro and colleagues (2020) also targeted the left TPJ in children and adolescents diagnosed with DD. They found anodal stimulation improved reading fluency, but specifically in the individuals with the most impaired fluency to begin with, suggesting that the proficiency level modulates stimulation effects. There has also been some success in improving fluency when targeting part of the visual system, left V5, in adults with DD fluent in Hebrew (Heth & Lavidor, 2015). After five anodal stimulation sessions over the course of two weeks, DD adults improved on RAN letters and numbers along with demonstrating an increase in the number of words read in one minute.

Though results tend to be mixed, there is an emerging trend that anodal tDCS can improve reading fluency, especially in DD readers. This tracks with evidence suggesting impaired readers show decreased reading network activation. The mixed results in typical readers may reflect the theory that typical readers already have a maximally efficient reading network, and therefore any manipulation or intervention to this system could decrease efficiency.

#### 4.4.2 Transcranial magnetic stimulation studies

Transcranial magnetic stimulation (TMS) is another neuromodulation method in which a high-intensity magnetic field is used to either facilitate or inhibit a brain region. TMS has been used for cortical mapping and clinical diagnoses (Hallett, 2007; Kobayashi & Pascual-Leone, 2003), but there are limited studies investigating TMS as an intervention to improve reading fluency. Most TMS studies have focused on how *disrupting* reading-related regions impacts different aspects of reading. Work has mostly concentrated on the dorsal visual stream responsible for reading-related eye movements and rapid visual attention, with TMS targeting posterior parietal regions (for review see Laycock & Crewther, 2008).

Only one study aimed to use TMS to improve reading fluency in readers with DD. Costanzo and colleagues (Costanzo et al., 2013) applied ten trains of 50 5-Hz TMS pulses to bilateral STG and IPL, two locations commonly implicated in reading, to native Italian speakers with DD. Mirroring the tDCS studies, targeting the left STG led to increases in single word reading speed.

### 4.5 Summary and discussion

Reading fluency research spanning multiple brain imaging modalities has identified fluency-related regions that align closely to the traditional reading network. Regions where GM is associated with fluency overlap extensively with fluency-related activation patterns, namely inferior frontal, temporoparietal, and occipitotemporal regions. We also see evidence that reading speed is associated with both functional and anatomical connections within the reading network.

The current review suggests two potential directions to investigate further: 1) subcortical structures and 2) WM tracts. First, subcortical structures have not been included in the traditional reading network but emerge consistently in fluency research (e.g., thalamus). Do these regions also support other aspects of reading, e.g., phonology or semantics, or are they specific to the speed of processing words? A meta-analysis of single word reading and lexical decision making in fMRI reported no subcortical structures (Murphy et al., 2019), though fMRI meta-analyses comparing typical and dyslexic readers across various reading tasks reported differences in the thalamus (Maisog et al., 2008; Richlan et al., 2009) and caudate (Richlan et al., 2009, 2011). These mixed results could be due to the inclusion of a wide variety of reading tasks (e.g., silent word reading, rhyme judgement, letter-sound integration) as well as the fact that fluency tasks were not widely included, if at all. Second, WM tracts associated with performance on timed tasks do not span the vast number of tracts associated with untimed reading tasks. The current review found converging evidence that reading fluency correlates with three WM tracts: left SLF, ILF, and SCR, which are only a subset of the WM tracts previously identified as associated with untimed word tasks.

One challenge of interpreting reading fluency research is that discrepancies in the direction of the relationship between performance and brain metrics start to emerge when comparing activation patterns from tasks and resting state FC changes following intervention in typical readers. Activation patterns and FC correlates of fluency tasks show a trend of greater activation and stronger FC in regions of the left hemisphere reading network that correspond with better fluency skills. Interestingly, interventions designed to increase reading speed in typical readers report faster reading related to *decreased* FC and *decreased* activity after training has been completed. A decrease in FC patterns and activity can be explained as the reading task becoming more automatic, transitioning into a background process requiring less activation, but this contradicts the idea that increases in activation and FC equates to faster reading. In impaired readers, interventions again support the idea that increased engagement of the reading network is associated with more efficient reading. When is an increase or decrease in activation more beneficial to reading speed and fluency, and in what populations? A clearer understanding of this will be important in designing neural-based approaches to improving reading fluency.

A second caveat stems from the use of region of interest (ROI) analyses. While this approach increases power for data analyses, it can ignore other brain regions that significantly contribute to reading fluency. ROI analyses tend to focus on cortical regions commonly implicated in the reading network, but whole-brain studies also suggest subcortical structures such as the cerebellum, thalamus, and caudate are involved in speeded reading. Given the well-documented roles of the basal ganglia (caudate) and cerebellum in procedural learning, these subcortical systems could provide critical support to the cortical reading network that enables reading processes to become faster and more efficient (e.g., Sokolov et al., 2017, Ullman 2016). Future research should explore the role these subcortical structures play in fluent reading.

Overall, studies investigating the neural bases of reading fluency make up a small portion of the reading literature, and we have yet to establish how to best distinguish the neural correlates of fluency from those of other reading domains. Understanding fluency at the neural level could help identify subtypes of reading-related disorders and assist in designing interventions for dysfluent individuals. It can also aid in differentiating fluency from phonological processing, which are often tapped by the same measures and share underlying processes. Understanding how regions outside of the reading network might be involved in fluent reading, including areas that support attention and processing speed, could aid more broadly in our understanding of how cognitive processes become efficient and automatic.

One way to parse out reading fluency measures is to look within the reading network at two circuits—the dorsal and ventral reading pathways (see **Figure 1**). The ventral pathway seems the intuitive choice to specifically support reading fluency because activation in these regions correspond with *rapid* identification of entire words (Kearns et al., 2019; Pugh et al., 2001). A second potential neural correlate of reading fluency is the cerebellum. The cerebellum has been consistently associated with both motor and cognitive skill automatization (Schmahmann et al., 2019; Sokolov et al., 2017), and this may manifest in the reading domain as fast, fluent reading. The right posterolateral cerebellum has also specifically been associated with reading, as it is functionally connected to the canonical left hemisphere reading network (Alvarez & Fiez, 2018) and has been associated with a variety of reading and reading-related tasks (King et al., 2019; Martin et al., 2015; Stoodley et al., 2012). While both the ventral pathway and cerebellum were identified as neural correlates of reading fluency in this qualitative review, a quantitative review of reading fluency neuroimaging studies may shed more light on whether these regions are consistently engaged during fluency tasks.

## 5. Activation likelihood estimate meta-analysis

To complement the qualitative literature review, we conducted an activation likelihood estimate (ALE) meta-analysis of fMRI studies of typical readers completing reading fluency tasks. We followed proposed guidelines for best practices when meta-analyzing neuroimaging studies (Müller et al., 2018) and PRISMA guidelines for reporting meta-analytic results (Page et al., 2021). We hypothesized that regions associated with the ventral reading pathway for rapid word recognition (i.e., OT cortex) would show converging activation across studies, along with the right posterolateral cerebellum which would help in the automatization, and therefore speed, of reading.

### 5.1 Methods

#### 5.1.1 Study selection

##### 5.1.1.1 Literature search

A systematic search for relevant neuroimaging publications was performed primarily using PubMed (https://pubmed.ncbi.nlm.nih.gov/), a life sciences and biomedical research database. Relevant publications were also identified through the sources for the qualitative review, three previously published reading neuroimaging meta-analyses (Barquero et al., 2014; Martin et al., 2015, 2016), and BrainMap’s literature repository via Sleuth 3.0.4 (www.brainmap.org/sleuth; Laird et al., 2011).

The PubMed literature search was completed on April 25, 2023. The search terms used were: “read*” AND (“speed” OR “fluency” OR “efficiency” OR “RAN” OR “rapid automatized naming“) AND (“fMRI” OR “neuroimaging”). This search resulted in 418 records. The BrainMap Sleuth literature search was completed on April 25, 2023. The search included papers that were categorized as having instructions that were “read.” This resulted in 135 records. Search terms were intentionally broad to encompass as many publications as possible because reading speed tasks can go by many names and descriptions.

##### 5.1.1.2 Inclusion criteria

All publications selected for the meta-analysis met the following criteria: 1) they used fMRI (PET and other functional imaging modalities were excluded), 2) they included results of a whole-brain analysis, 3) a reading fluency/speed task was completed (either inside or outside the scanner), 4) contrasts of interest were performed on typical readers, and 5) results were reported in standard coordinate space (e.g., MNI, Talairach). Each abstract and/or full text was assessed for exclusion in the following order (**Figure 2**): review articles, meta-analyses, case studies, study did not include fMRI results, study did not include a typical reader group or separate data analyses of only the typical reader group, no whole-brain analyses were conducted, no relevant speed-related reading task or relevant analyses were not conducted on speed-related aspect of reading, and miscellaneous (e.g., subjects were not human, study’s purpose was to review or critique fMRI analysis methods).

**Figure 2.**
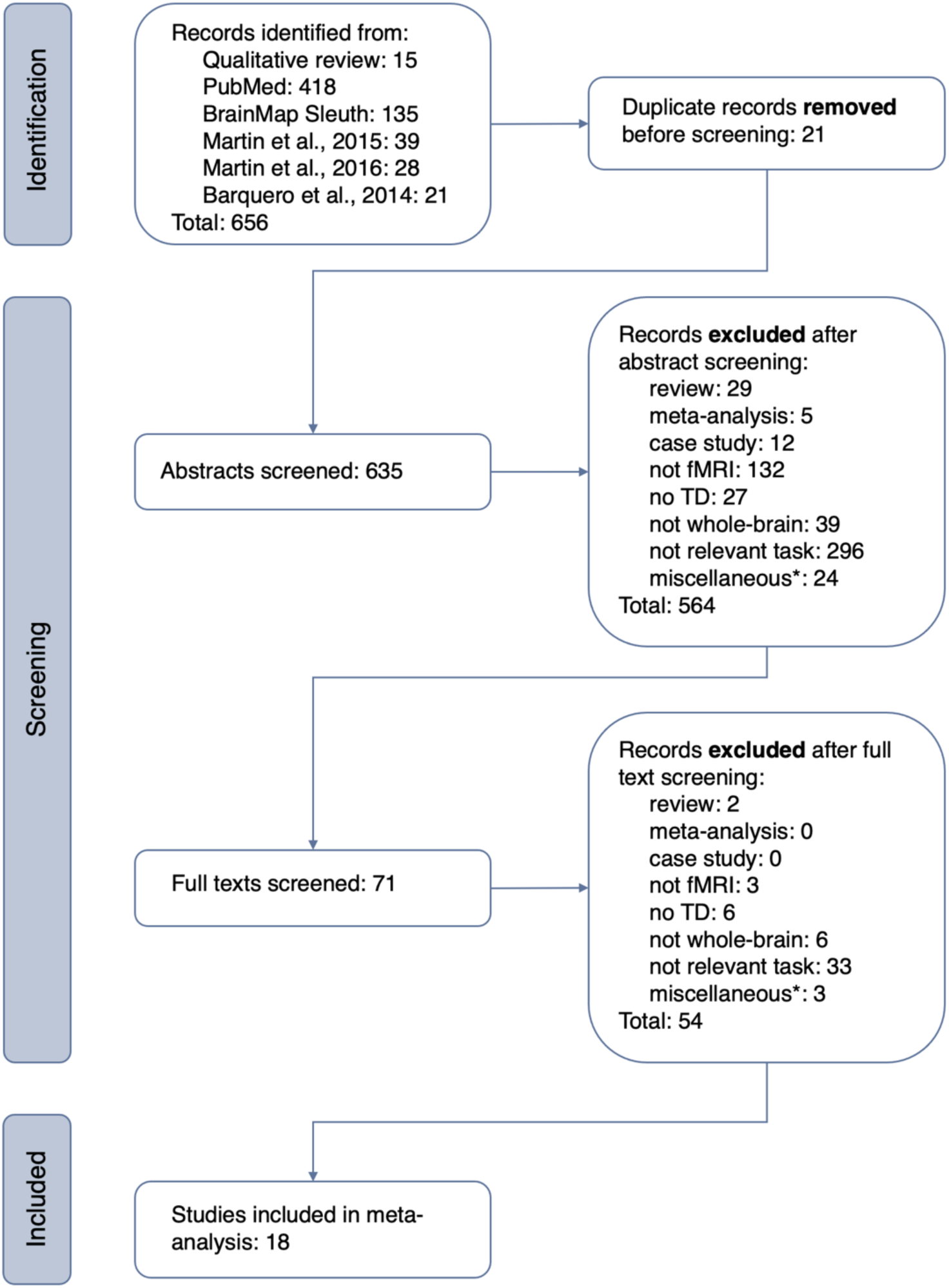
PRISMA flowchart. *Miscellaneous reasons for exclusion included the following: study was reported as *under review* and a published version was not found; study included a reading speed task that did not include a relevant contrast evaluating speed; relevant data was repeated in another publication and that publication was retained in the meta-analysis; non-human subjects; methods paper

**Figure 3.**
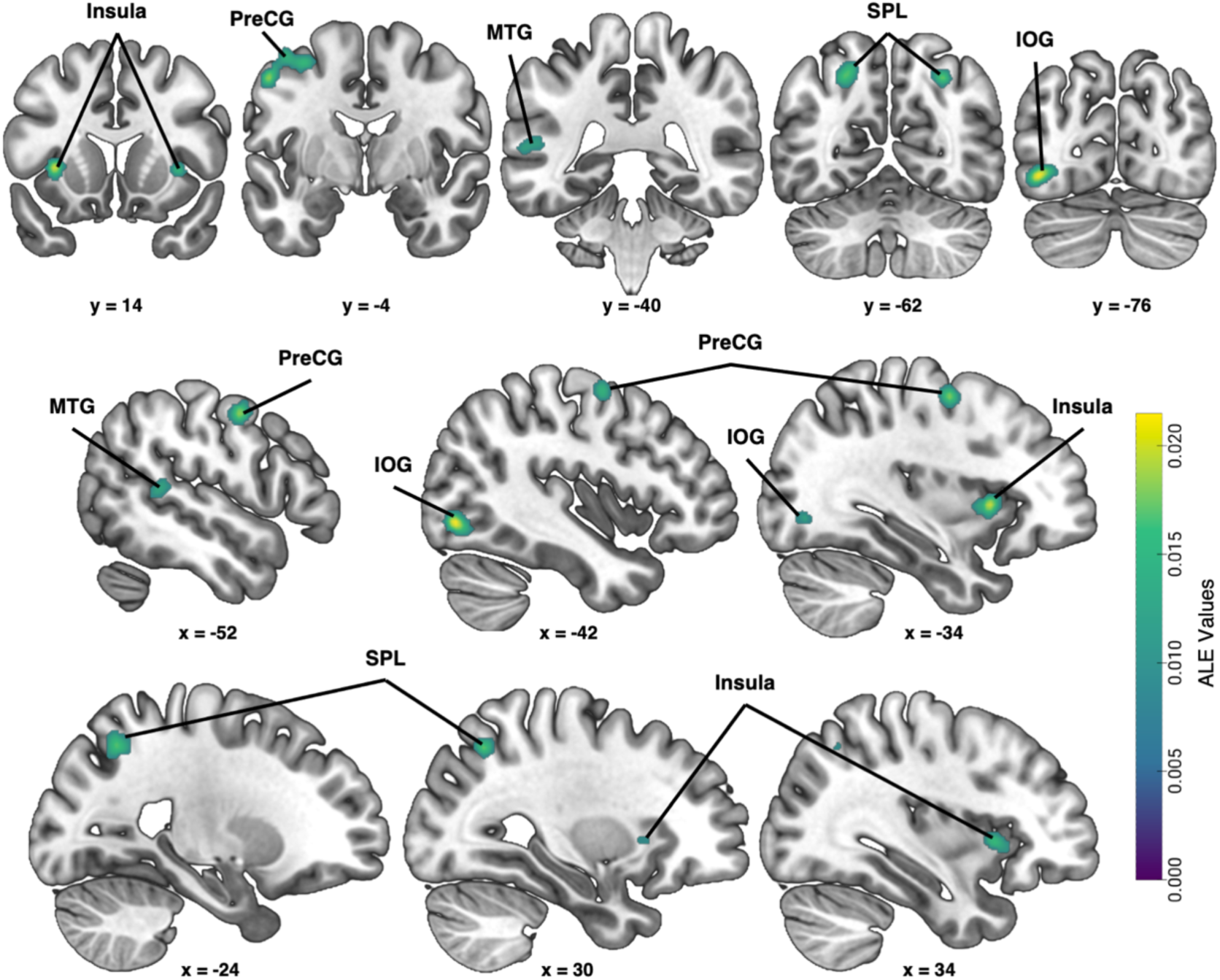
Areas of convergence identified by ALE meta-analysis, voxel p_uncorr_ < 0.001, cluster p_FWE_ < 0.05. Abbreviations: IOG = inferior occipital gyrus, MTG = middle temporal gyrus, PreCG = precentral gyrus, SPL = superior parietal lobule.

#### 5.1.2 Data extraction and contrasts of interest

First, each publication’s abstract was evaluated based on the inclusion/exclusion criteria. If a decision could not be made based on the abstract, we performed a full text screening so more information could be gathered. All publications that passed abstract screening were further screened at the full text level.

Data were extracted from the main publications and supplemental materials. Information describing xyz coordinate results from the contrasts of interest, statistical software and thresholds, sample size, and relevant demographic information about the participants were collected.

To examine the neural correlates of reading fluency, contrasts were chosen that reflected increased activation associated with increased fluency or speed of reading (e.g., RAN letters > fixation, accelerated > normal > constrained reading rates). Because only two of the 18 final publications reported decreased activation associated with increased reading fluency, and additionally because only one contrast per publication was selected to avoid within-group effects (Müller et al., 2018; Turkeltaub et al., 2012), we decided to only analyze positive relationships. When multiple fluency tasks were administered, the task most similar to real-world reading was chosen (e.g., RAN letters was chosen over RAN digits or objects, sentences were selected over single word reading).

After screening for the appropriate inclusion criteria and contrasts of interest, 18 publications (**Figure 2**, **Table 1, Table S1**) met inclusion criteria for the meta-analysis, which is consistent with the minimum number recommended in neuroimaging meta-analysis guidelines (17-20; Eickhoff et al., 2016; Müller et al., 2018).

**Table 1.** List of reading fluency studies included in the ALE meta-analysis.

|  | Author | Year | N | Task Type | Response Modality | Task inside or outside scanner | Contrasts of Interest | # Foci |
| --- | --- | --- | --- | --- | --- | --- | --- | --- |
| 1 | Barbeau et al. | 2017 | 14 | paragraph reading | overt reading | outside | positive association between MR signal and faster paragraph reading | 1 |
| 2 | Benjamin & Gaab | 2012 | 13 | fluent sentence reading under normal, constrained, or accelerated presentation rates | covert reading; button press (indicate start and stop) | inside | accelerated > rest | 4 |
| 3 | Binder et al. | 2005 | 24 | single word and nonword reading | overt reading | inside | activations correlated with RT (continuous variable) | 43 |
| 4 | Buchweitz et al. | 2014 | 11 | sentence reading (familiar and unfamiliar topics) under normal and fast presentation rates | covert reading; button press (confirming sentence comprehension) | inside | speed reading > normal reading | 4 |
| 5 | Christodoulou et al. | 2014 | 12 | sentence reading under slow, medium, and fast presentation rates | covert reading; button press (indicate if sentence semantically plausible) | inside | fast > medium > slow | 12 |
| 6 | Cummine et al. | 2015 | 15 | RAN letters, rapid word reading | covert reading; button press (indicate end of card) | inside | conjunction of RAN letters and words | 11 |
| 7 | Dahhan et al. | 2020 | 18 | letter and object naming speed | overt reading | inside | Letter and Object NS > fixation | 22 |
| 8 | Fu et al. | 2002 | 7 | single word reading under slow and fast presentation rates | covert reading | inside | fast > fixation (pinyin) | 39 |
| 9 | Hashimoto & Sakai | 2003 | 18 | sentence reading under slow, normal, fast presentation rates | overt reading | inside | fast > normal | 3 |
| 10 | Langer et al. | 2015 | 15 | fluent sentence reading under normal, constrained, and accelerated presentation rates | covert reading; button press (indicate start and stop) | inside | accelerated > rest (sentences) | 5 |
| 11 | Mechelli et al. | 2000 | 6 | single word reading under 20, 40, and 60 words per minute presentation rates | covert reading | inside | positive linear effect (60wpm > 40wpm > 20wpm) | 9 |
| 12 | Misra et al. | 2004 | 12 | RAN letters | covert reading; button press (indicate when end of line was reached) | inside | letter naming > fixation | 14 |
| 13 | Miura et al. | 2003 | 23 | sentence reading | covert reading | inside | positive correlation between MR signal (meaningful sentence reading) and reading speed | 6 |
| 14 | Ozernov-Palchik et al. | 2022 | 26 | sentence reading under constrained, normal, and acceleration presentation rates | covert reading; button press (confirming sentence comprehension) | inside | accelerated > rest (Time 2) | 2 |
| 15 | Qian et al. | 2015 | 20 | RAN digits; coherent motion detection paradigm | covert naming; button press (indicate motion direction) | outside | positive correlation between RAN and brain activity associated with coherent motion task | 9 |
| 16 | Turkeltaub et al. | 2003 | 26 | RAN letters | overt reading | outside | positive correlation between RAN letters and brain activation | 8 |
| 17 | Welcome & Joanisse | 2012 | 20 | TOWRE-2 Sight Word Efficiency (SWE); word reading task | overt reading; button press (indicate case, phonology, or semantic decisions) | outside | positive correlation between SWE and brain activation during word reading | 3 |
| 18 | Zou et al. | 2016 | 21 | RAN letters | covert reading; button press (indicate when end of line was reached) | inside | RAN letter > null | 6 |

*Note:* Each row indicates one study and one contrast of interest. N indicates the sample size. Number of foci were the number of foci (x, y, z coordinates) from the contrast of interest that were entered into the meta-analysis.

#### 5.1.3 Anatomic likelihood estimate (ALE) meta-analysis procedure

The meta-analysis was performed using BrainMap’s GingerALE 3.0.2 (www.brainmap.org/ale; Eickhoff et al., 2009, 2012; Laird et al., 2005), a method originally described by Turkeltaub et al. (Turkeltaub et al., 2002), which identifies neural areas of convergence across experiments based on lists of peak activation foci (e.g., xyz coordinates) extracted from published studies. The current version of GingerALE incorporates random effects and adjusts the full-width half-maximum (FWHM) of the Gaussian function depending on the sample size of each study, such that studies with larger sample sizes are blurred with a smaller FWHM and coordinates from studies with smaller sample sizes are blurred with a larger FWHM.

All extracted foci loaded into the analysis were in MNI space. If a study reported foci in Talairach space, the ‘Convert Foci’ tool from GingerALE was used to convert into MNI space using the icbm2tal transform (Lancaster et al., 2007). This conversion method is recommended over the Brett transform as it has better fit and improves the overall accuracy of the meta-analysis (Laird et al., 2010).

The ALE map was thresholded using an uncorrected voxel-level threshold of p < 0.001 and FWE-corrected cluster-level threshold of p < 0.05 (Eickhoff et al., 2012, 2016).

Results were visualized with MRIcroGL overlaid on the ch2better brain template. Anatomical labels were assigned to each cluster from the Talairach Daemon (Lancaster et al., 1997, 2000), which is automatically included in the GingerALE meta-analysis output.

#### 5.1.4 Comparisons with Neurosynth database

To determine if the regions showing converging activation across reading fluency studies are generally associated with reading or reading-related behaviors (e.g., reading, words, syntax, language, etc.), the peak coordinates identified by the meta-analysis were entered into the Neurosynth data base (https://neurosynth.org; Yarkoni et al., 2011), an online repository of more than 14,000 studies reporting data from over 150,000 brain regions. Neurosynth also includes a database of >1300 terms extracted from study abstracts which can be associated with the corresponding studies’ brain coordinates. This allowed us to evaluate whether reading fluency was associated with the reading network specifically (associated terms mostly include reading and/or language) or more domain-general regions supporting cognitive processing. Associated terms were included if their z-score was greater than 4. Z-scores were automatically calculated by Neurosynth via two-way ANOVAs testing for a non-zero association between terms and voxel activation. This provided an indication of how consistently activation in a specific brain region occurs for studies reporting a specific term versus studies not reporting the term.

### 5.2 Results

Figure 3 and **Table 2** show brain regions where increased activation was consistently associated with reading fluency / RAN measures as well as the top Neurosynth associations with the peak ALE coordinates. The areas of convergence found in this analysis overlap extensively with the qualitative review of reading fluency literature and the traditional cortical reading network, including the occipitotemporal (OT) cortex, left middle temporal gyrus (MTG), left superior parietal lobule (SPL), and left precentral gyrus. Additional cluster information can be found in **Table S2**. Except for the bilateral insula, each region was associated with reading-related terms from the Neurosynth repository.

**Table 2.** Results of the ALE meta-analysis.

| Cluster | Anatomical Label | Volume (mm<sup>3</sup>) | x | y | z | ALE Value | Articles with Foci | Neurosynth Top Associations* |
| --- | --- | --- | --- | --- | --- | --- | --- | --- |
| 1 | Precentral Gyrus | 2136 | -52 | -4 | 46 | 0.0187 | 2, 6, 7, 8, 12, 13, 18 | language, motor, **speech**, **phonological**, listening, **reading**, auditory, movements, production, **words**, rehearsal, vocal, sounds, **spoken**, articulatory, eye movement, **pseudowords**, covert, sequences, **word**, eye movements, listened, motor imagery, **linguistic**, eye, musical, acoustic, tasks, **sentences**, movement, pitch, saccades |
|  | Precentral Gyrus |  | -34 | -6 | 54 | 0.0184 |  |  |
|  | Precentral Gyrus |  | -44 | -4 | 58 | 0.0152 |  |  |
| 2 | Inferior Occipital Gyrus | 1520 | -42 | -76 | -6 | 0.0215 | 3, 4, 5, 11, 15, 18 | visual, objects, **reading**, motion, **orthographic**, stream, face recognition, object, visual motion, selective, naming, faces, visual stream, **read**, **word**, **readers**, recognize, color, **words** |
| 3 | Superior Parietal Lobule | 1304 | -24 | -62 | 48 | 0.0172 | 3, 7, 8, 12, 15, 18 | visual, task, tasks, attentional, **English**, working memory, gain, working, interference, **reading**, **word**, **orthographic**, demands, **phonological**, attentional control, distractors, visual attention, attention, **Chinese**, color, rotation, load, **languages**, memory, arithmetic, **visual word**, orienting, shifting |
| 4 | Superior Parietal Lobule | 1024 | 30 | -62 | 46 | 0.0176 | 3, 4, 7, 8 | calculation, task, arithmetic, working memory, working, tasks, **reading**, visually, mood, visual, **Chinese**, symbolic, load, rotation, visually presented, **characters**, memory |
|  | Precuneus |  | 22 | 066 | 50 | 0.0160 |  |  |
| 5 | Middle Temporal Gyrus | 984 | -50 | -44 | 8 | 0.0152 | 5, 7, 8, 10, 18 | **sentences**, **language**, **reading**, **spoken**, **words**, **sentence**, **syntactic**, **speech**, listening, auditory, comprehension, **linguistic**, **visual word**, **word**, **word form**, **lexical**, **English**, **phonological**, **verbs**, unimodal, **pseudowords**, **written**, **languages**, **orthographic**, **read**, vocal, audiovisual, speakers, acoustic, **dyslexia**, **sentence comprehension**, **semantic**, conflicting, visual auditory, auditory visual, **Chinese**, **verb**, **characters**, face, **readers** |
|  | Middle Temporal Gyrus |  | -60 | -38 | 8 | 0.0139 |  |  |
| 6 | Insula | 888 | -34 | 14 | 2 | 0.0203 | 3, 7, 9, 12 | pain, painful, secondary somatosensory, sensation, noxious |
| 7 | Insula | 640 | 34 | 16 | 0 | 0.0165 | 2, 3, 7 | pain, gain, painful, response inhibition, load, task, stop, stop signal, taste, behavior, signal task, secondary somatosensory, PTSD, calculation |

\*Top associations were included if $z > 4$ . Associations are listed in order of greatest z-score to lowest z-score. Only associations that did not include the name of the brain region itself (e.g., temporal, fusiform) are listed. Only peak coordinates for each cluster identified in the ALE meta-analysis were used.

### 5.3 Discussion

The aim of this meta-analysis was to determine the converging neural correlates of reading fluency in typical readers. We hypothesized that convergence would be seen in the left-lateralized ventral reading pathway and right posterolateral cerebellum to account for rapid word recognition and automatized reading. Eighteen studies met criteria for inclusion in the ALE meta-analysis. From this, we identified seven clusters associated with reading fluency across studies: left precentral gyrus, left inferior occipital gyrus (IOG), bilateral superior parietal lobule (SPL), left middle temporal gyrus (MTG), and bilateral insula. Our hypothesis was partially supported, as the ventral reading pathway OT region was consistently recruited for reading fluency tasks, although these tasks also engaged dorsal reading pathway regions such as the precentral gyrus. Our analysis identified both canonical components of the reading network (left OT, left MTG) as well as regions more broadly associated with cognitive task performance (SPL, insula). The Neurosynth associations for the peak coordinates are consistent with the interpretation that fluent reading taps not only the core reading network but also recruit brain areas that are necessary for cognitively demanding tasks.

#### 5.3.1 Ventral reading pathway is engaged during rapid reading

The ventral reading pathway, composed of left OT regions (e.g., IOG, fusiform gyrus, inferior temporal gyrus) and the MTG, is recruited when typical readers rapidly recognize familiar words (Kearns et al., 2019; Pugh et al., 2001). The left IOG was one of the largest clusters identified in the meta-analysis, and a portion of the cluster overlapped with the fusiform gyrus (see **Table S2**). Left IOG activation has been an area of convergence in multiple meta-analyses of reading (Houdé et al., 2010; Martin et al., 2015; Murphy et al., 2019). This area tends to be more sensitive to meaningful words rather than scrambled words as seen in Chinese characters (Zhang et al., 2018), similar to the visual word form area (VWFA) also found in the OT region, though more anteriorly. Activation in this region occurs very quickly during reading, with activation in the left fusiform gyrus approximately 150 to 200ms after seeing a word (Cohen et al., 2000). This early activation in the fusiform is associated with the ability to determine if the visual stimulus is a linguistic stimulus (e.g., a word or pseudoword) rather than a non-linguistic stimulus (Cohen et al., 2000; Pugh et al., 2001). The early recognition of linguistic stimuli can assist fluent reading, and neuroimaging research supports this. Six of the studies identified in the current meta-analysis reported IOG activation during a reading fluency task. Binder and colleagues (2005) found increased IOG activation with faster response times when reading a list of words, regardless of whether they were real words, irregular words, or nonwords. A similar effect is evident with RAN letters: greater fusiform activation occurred in response to rapidly reading a matrix of single digits or letters (Qian et al.,2015; Zou et al., 2016). Last, fusiform activation levels showed a gradient effect when examining multiple speeds of reading (e.g., slow, normal, fast). As reading speed increased, so did fusiform activation (Buchweitz et al., 2014; Christodoulou et al., 2014; Mechelli et al., 2000).

Outside of the studies included in the meta-analysis, the IOG and fusiform are consistently associated with reading fluency. Left IOG GM volume is correlated with reading speed in children regardless of their age, with less GM associated with better reading speed (Simon et al., 2018). In pre-readers, fusiform activation predicts later fluency, as measured through RAN (Liebig et al., 2021). There is also evidence of functional and structural differences in the fusiform in dysfluent readers, such as those diagnosed with DD. These readers tended to have less fusiform gray matter (Martins et al., 2021) and hypoactivity (Richlan et al., 2011) compared to their more fluent counterparts. Similar patterns of hypoactivity can also be seen in the left IOG of readers with DD (Zhang & Peng, 2022). Additionally, the correlations between fusiform activation and reading fluency in typical readers were not evident in those with a diagnosis of developmental dyslexia (Hashimoto et al., 2020).

The left MTG was another region of convergence in the current meta-analysis that is also considered part of the ventral reading pathway. The left MTG tends to be associated with extracting word meaning rather than quickly identifying linguistic stimuli, though there is also evidence for its involvement in morphological processing (Devlin et al., 2004; Fruchter & Marantz, 2015), the ability to understand the basic components that make up a complex word (such as “jumped” being composed of the verb “jump” and “ed” to indicate past tense; Levesque et al., 2021). Being able to quickly parse parts of a word to understand its meaning is also essential to fluent reading. Five studies included in the meta-analysis reported activation in the MTG. Dahhan et al. (2020) and Qian et al. (2015) found that faster automatic naming speed correlated with increased MTG activation, and MTG activation also increased as word (Fu et al., 2002) or sentence (Christodoulou et al., 2014; Langer et al., 2015) presentation rates increased.

These results support the hypothesis that the ventral reading pathway contributes to reading fluency. Activation in these regions increases as reading speed increases in single letter, single word, and sentence reading, regardless of whether the speed is controlled by the study paradigm or calculated as reaction times from the participant.

#### 5.3.2 Precentral gyrus as a dorsal reading pathway node or domain-general support

The dorsal reading pathway is traditionally composed of left STG, AG, SMG, and precentral gyrus (Kearns et al., 2019). This pathway responds more to unfamiliar words or words that require a sound-it-out approach to decoding (Kearns et al., 2019; Pugh et al., 2001). While we did not hypothesize that reading fluency would specifically tap the dorsal pathway, our meta-analysis identified the left precentral gyrus as a region of convergence across fluency studies.

Activation patterns of the precentral gyrus have been correlated with many aspects of reading. There is evidence showing its role in the motor aspects of language and reading, such as articulation (Kearns et al., 2019; Silva et al., 2022) and saccadic eye movements during reading (Zhou et al., 2016). The current precentral gyrus findings overlap with the mid-precentral gyrus, a region that may support speech motor planning of phonological information (Silva et al., 2022). Nonmotor reading contributions of the precentral gyrus point to phonological processing (Kaestner et al., 2021; MacSweeney et al., 2009; Sakurai et al., 2018; Yen et al., 2019) and assistance in letter identification through communication with the fusiform gyrus very early in word processing (Kaestner et al., 2021). More generally, the left precentral gyrus might also contribute to executive function processes that are required during fluent reading (K. Wang et al., 2020). Associated terms from Neurosynth do not clear up this divide between reading-specific and domain-general contributions as the list consists of both types of terms, nor even more specifically with how this region supports motor aspects of reading (i.e., through eye movements, mouth movements, or subvocal articulation/planning).

Seven studies included in the current meta-analysis contributed to the left precentral gyrus clusters. Four of those studies found positive correlations between precentral gyrus activation and RAN speed (Cummine et al., 2015; Dahhan et al., 2020; Misra et al., 2004, Zou et al., 2016). Precentral gyrus activation was also positively correlated with faster single word reading (Fu et al., 2002) and sentence reading (Benjamin & Gaab, 2012; Miura et al., 2003). The current meta-analysis cannot determine if there was an effect of covert versus overt reading during the task, and therefore a motor effect, as only six of the 18 studies included used overt reading, though only one of the six studies contributed to the precentral gyrus cluster found in the meta-analysis.

Unfortunately, we cannot say with certainty how the left precentral gyrus contributes to reading based on the current results. The cluster overlaps with reading-related associations and with eye and mouth movements via Neurosynth. Additionally, the meta-analysis identified convergence in the left fusiform gyrus, part of the left IOG cluster, which may communicate with the precentral gyrus when identifying letters (Kaestner et al., 2021). We suggest the left precentral gyrus supports both motor and nonmotor aspects of reading in a regionally-specific manner (K. Wang et al., 2020).

#### 5.3.3 Integration of ventral and dorsal pathways during fluent reading

The current meta-analysis revealed both ventral and dorsal reading pathway regions contributing to reading fluency, suggesting that fluent reading requires the successful integration of these two pathways. Blomert (2011) reviewed the literature on letter-sound integration in typical and dyslexic readers and suggested a parietal-temporal-occipital network that automatized letter-sound pairings. To achieve this orthographic-phonological skill, superior temporal and parietal regions support sound processing while occipitotemporal areas identify letters. The superior temporal sulcus then integrates these two modalities. Event-related potential (ERP) data from a letter-sound integration automation paradigm supports the notion that skilled readers perform letter-sound integration very quickly, within approximately 100-200ms, whereas poor readers and those diagnosed with dyslexia did not show automation of this skill. Blomert argues that automaticity of letter-sound integration can lead to fast, fluent reading, and the lack of automaticity in those diagnosed with dyslexia, along with decreased fusiform response to letters, can result in slow, dysfluent reading.

Additional support for dorsal and ventral reading pathways working together during fluent reading comes from RAN neuroimaging studies. Qian and colleagues (Qian et al., 2016) collected resting state functional connectivity data from typical readers who completed a RAN task outside of the scanner. RAN scores were associated with functional connectivity in right SPL, left MTG, and left fusiform areas, with stronger connectivity correlated with faster naming. Further examination of RAN activation patterns also suggests that rapidly naming alphanumeric stimuli, words, and nonwords recruits both ventral (middle occipital gyrus, lingual gyrus) and dorsal (precentral gyrus, posterior parietal cortex) pathways (Cummine et al., 2015).

These studies support the view that the dorsal and ventral reading pathways must work together during timed reading tasks. The integration of these two pathways may support fast integration of phonological and orthographic characteristics of words, which in turn leads to fluent reading.

#### 5.3.4 Superior parietal lobule and insular contributions to reading

##### 5.3.4.1 Superior parietal lobule

The term “posterior parietal cortex” has been used to describe parietal regions associated with the reading network, but it is specifically the *inferior* parietal lobule that is most commonly associated with reading, rather than the SPL. Regardless, bilateral SPL clusters extending into the right precuneus were found as regions of convergence in the current meta-analysis. In a large meta-analysis of typical readers, Martin and colleagues (Martin et al., 2015) found left SPL activation was associated with reading in children, but this cluster was not found in adult readers. Based on these findings, how might this area be involved in reading?

One possibility comes from the attention literature. The bilateral SPL is considered part of the orienting network, a top-down visuospatial attention network composed of the intraparietal sulcus, SPL, and frontal eye fields which help guide attention by directing one to the necessary and appropriate sensory information during a task (Petersen & Posner, 2012). Bilateral SPL is also associated with visual attention span, which has been suggested to contribute to developmental dyslexia (Bosse et al., 2007). Typical readers tend to recruit bilateral SPL during tasks designed to assess visual attention, whereas those with DD either do not engage the SPL or show decreased activation in response to higher visual attention loads. For example, when single letters are “flanked” by additional letters to either side, therefore increasing the amount of noise around the target stimuli and requiring more targeted attention, typical readers show increased activation in bilateral SPL compared to when no flankers are present (Peyrin et al., 2011). Individuals with DD did not show this pattern. Bilateral SPL is also recruited in typical reading development when words become more visually degraded and harder to read, also arguably requiring more targeted attention (Cohen et al., 2008). Last, Reilhac and colleagues (Reilhac et al., 2013) suggest the SPL is used for letter detection. Their participants completed a letter identity task in which they had to determine if two letter strings presented one after another were the same or different. While the authors argue that the SPL activation was associated with letter identification (as compared to a control task where the participant determined if a frame was presented around the letter strings or not), the task itself could still recruit the orienting network due to the higher attentional load during the letter string condition.

In the current meta-analysis, three studies reported bilateral SPL clusters, one study reported only the right SPL, and three studies only the left SPL. Faster response times when reading words, irregular words, and nonwords (Binder et al., 2005), faster rapid naming speed for letters and objects (Dahhan et al., 2020), and faster single word reading (Fu et al., 2002) recruited bilateral SPL. Interestingly, three other rapid naming studies (letters and digits) reported only left SPL activation (Misra et al., 2004; Qian et al., 2015; Zou et al., 2016) while faster sentence reading recruited only the right SPL (Buchweitz et al., 2014). While there is no obvious reason for this differential hemispheric recruitment, a working theory could be that the recruitment of the SPL is related to the amount of cognitive load required to perform the task. Simply rapidly naming single letters is quite easy for a typical reader, whereas the other tasks engaging the right SPL were more complex and arguably placed higher demands on attention.

##### 5.3.4.2 Insula

The bilateral insulae were additional areas of convergence in the current meta-analysis that are not commonly discussed as part of the neurobiological underpinnings of reading. They were also the only regions that were not associated with reading-related keywords as per the Neurosynth top associations. Most studies from the Neurosynth database reporting results in the insula focused on pain and response inhibition.

In general, the insula has been associated with a range of cognitive abilities (Shelley & Trimble, 2004), and more recently has been functionally parcellated into three subdivisions (Chang et al., 2013). The current meta-analytic insular clusters overlap with the anterior dorsal insula, which is associated with higher-order cognition and executive functions as identified through forward and reverse inference using the Neurosynth database. This insular subdivision was overwhelming “cognitive” compared to the other two divisions, which were mostly associated with either emotion, gustation, and olfaction processing or pain, somatosensory, and sensorimotor processing.

While there is a lack of discussion of the insula in the reading literature, it has been reported as engaged during reading tasks in multiple meta-analyses. In a meta-analysis studying the neural correlates of typical reading, the left anterior insula was part of a large left inferior frontal gyrus cluster (Martin et al., 2015). In meta-analyses focusing on studies comparing typical readers to those diagnosed with developmental dyslexia, there was a consistent pattern of hypoactivation of the right insula (Maisog et al., 2008) and hyperactivation of the left insula in the dyslexia group compared to those with typical reading development (Martin et al., 2016; Richlan et al., 2009), though the latter cluster may be driven by languages with deep orthographies (Richlan et al., 2009).

In the current meta-analysis, five studies contributed to the insula clusters. Two studies used rapid naming paradigms and associated faster rapid naming with increased insula activation (Dahhan et al., 2020; Misra et al., 2004). Faster single word reading and faster sentence reading also correlated with increased insula activation (Benjamin & Gaab, 2012; Binder et al., 2005; Hashimoto & Sakai, 2003).

Additional neuroimaging studies also point to a connection between the insula and reading fluency. Gray matter volume of the right insula positively correlated with reading fluency scores (He et al., 2013), while volume in the left insula was negatively associated with rapid naming of pictures (Jednoróg et al., 2015). These insula clusters overlap with the dorsal and ventral anterior subdivisions identified by Chang and colleagues (Chang et al., 2013), which support initiating and sustaining attention and error detection (Droutman et al., 2015). These behaviors are important during reading—proper attention must be allocated to the task, and errors in reading must be identified for accurate reading to occur. Therefore, similar to the SPL, the insular cortex may provide domain-general support to fluent reading.

### 5.4 Limitations

The current meta-analysis results should be considered in the light of a few limitations. Some studies included in the meta-analysis had small sample sizes, which lack appropriate statistical power (Lin, 2018; Turner et al., 2013). Because we identified only 18 studies that fit the inclusion criteria, we decided to include all relevant papers regardless of sample size, with the intention to include more studies as they are published. The purpose of these results is to establish a starting point for future research investigating the neural underpinnings of reading fluency.

It is also important to determine if the neural activation that is measured as reading speed increases is actually due to increased reading fluency or to increased cognitive load, because larger amounts of stimuli are presented in roughly the same amount of time. While this is difficult to disentangle, we argue that the contrasts of interest entered into the meta-analysis assessed reading speed specifically, as not all of the tasks fit the idea that increased speed equates to higher cognitive load. The RAN tasks, which were contrasted with a rest condition, arguably add less cognitive load compared to studies controlling the speed of sentence presentation rates, especially during accelerated presentation rate conditions. The amount of cognitive load required for RAN can be considered relatively low because the purpose of the task is to use stimuli that are highly familiar and automatically recalled. The types of tasks contributing to each neural region of convergence in the current meta-analysis were composed of both low and high cognitive load studies, i.e., studies using RAN tasks (lower cognitive load) and studies using whole sentences or varying presentation rates (higher cognitive load), which suggests cognitive load may not be a driving factor. That said, performing any task at speed will tap additional neural resources relative to untimed measures.

### 5.5 Conclusions

The goal of the current meta-analysis was to assess areas associated with reading fluency across published studies. Convergence was found in the traditional reading network, spanning both the dorsal and ventral reading pathways, though most studies reported robust engagement of the ventral pathway. This suggests that the ventral pathway is heavily recruited for automatic, fluent reading. Additional regions outside the traditional reading network are also recruited during fluent reading and RAN tasks, most likely supporting attention and error-monitoring. These results are consistent with the neural regions predicted from our qualitative review of the literature.

## 6. General conclusions

The qualitative review of reading fluency across the neuroimaging literature suggests fluent reading recruits nearly all areas associated with the cortical reading network, along with subcortical and cerebellar regions. The quantitative meta-analysis examining convergence across fMRI studies suggested that rapid reading and RAN measures primarily tap the ventral reading pathway and outside-network areas supporting attention. Both these approaches indicated that the neuroimaging literature of reading fluency is quite small compared to other aspects of reading.

Additional work must be done to elucidate how reading fluency is processed in typical reading development and then expanded upon in reading disability. We have provided support for the relationship between rapid reading and the ventral reading pathway, which may be a starting point for the future of this area of study. These results, in turn, may also aid in understanding different types of reading disability as viewed through the double-deficit hypothesis (e.g., Norton et al., 2014; Norton & Wolf, 2012; Wolf & Bowers, 1999). The ultimate goal of understanding the neural correlates of fluent reading is to identify biomarkers for successful reading interventions and potential targets for approaches to improve reading speed in reading disabled individuals.

---

**Table S1.** Studies included in the meta-analysis.

| Author | Year | N | Age Group | % Female | Mean Age (yrs) | Age SD (yrs) | Age Range (yrs) | Standard Space | Software | Critical Threshold | Corrected or Uncorrected |
| --- | --- | --- | --- | --- | --- | --- | --- | --- | --- | --- | --- |
| Barbeau et al. | 2017 | 14 | adults | NA | 24.9 | 3.7 | NA | MNI | SPM8 | $p < 0.005$ | cluster-level FDR-corrected $p < 0.05$ |
| Benjamin & Gaab | 2012 | 13 | adults | 92 | 24.05 | 4.48 | 19.16-33.24 | MNI152 | FSL | $z > 2.3$ | cluster corrected $p < 0.05$ |
| Binder et al. | 2005 | 24 | adults | 50 | 27.5 | NA | 18-48 | Talairach | AFNI | voxel-wise $p < 0.001$<br>($t \geq 3.76$) | cluster corrected $p < 0.05$ |
| Buchweitz et al. | 2014 | 11 | adults | 36 | 21.64 | 2.54 | 18-27 | MNI | SPM2 | voxel-wise $p < 0.001$ | $k > 20$ |
| Christodoulou et al. | 2014 | 12 | adults | 42 | 22.5 | 3.1 | 18-28 | MNI | SPM8 | $p < 0.001$ | cluster-level FDR corrected, $k > 10$ |
| Cummine et al. | 2015 | 15 | adults | 33 | 20.63 | 2.67 | 18-26 | MNI | SPM8 | voxel-wise $p < 0.001$ | cluster-level $p < 0.001$ |
| Dahhan et al. | 2019 | 18 | adults | 89 | 24.1 | 1.89 | 21-26 | Talairach | BrainVoyager | $p < 0.01$; $t = 2.90$ | cluster corrected $p < 0.05$ |
| Fu et al. | 2002 | 7 | adults | NA | NA | NA | 25-38 | Talairach | FMRIB Easy Analysis Tool | $Z > 3.1$ | cluster corrected $p < 0.01$ |
| Hashimoto & Sakai* | 2003 | 15 | adults | NA | NA | NA | NA | MNI | SPM99 | voxel-level $p < 0.05$ | "corrected for multiple comparisons" |
| Langer et al. | 2015 | 15 | children | 47 | 9.9 | 1.55 | 8.3-12.5 | MNI152 | FSL | $Z > 2.3$ | cluster corrected $p < 0.05$ |
| Mechelli et al. | 2000 | 6 | adults | 17 | 24 | NA | 20-34 | Talairach | SPM99 | $p < 0.05$ | corrected for multiple comparisons; $k > 5$ |
| Misra et al. | 2004 | 12 | adults | 83 | NA | NA | 18-25 | MNI305 | SPM99 | $p < 0.001$ | uncorrected for multiple comparisons; $k > 10$ |
| Miura et al. | 2003 | 23 | adults | 30 | 23 | NA | 18-44 | MNI | SPM99 | $p < 0.05$ | corrected for multiple comparisons |
| Ozernov-Palchik et al. | 2022 | 26 | children | 42 | 9.5 | 1.17 | NA | MNI152 | FSL | $z = 2.3$ | cluster corrected $p < 0.05$ |
| Qian et al. | 2015 | 20 | adults | 55 | 22.5 | 1.59 | 20-24 | MNI | SPM8 | $p < 0.05$ | FDR corrected and $k > 5$ |
| Turkeltaub et al. | 2003 | 41 | children | 54 | 14.1 | 5.6 | 7.0-21.5 | Talairach | SPM99 | $p < 0.0001$ | $k > 25$ at $p < 0.001$ |
| Welcome & Joanisse | 2012 | 20 | adults | 65 | 29.7 | 13.7 | 19-59 | Talairach | AFNI | $p < 0.001$ | $k > 243$ |
| Zou et al. | 2016 | 21 | children | 33 | 12.3 | 0.6 | 6-25 | MNI | SPM5/SPM8 | $p < 0.05$ | FWE corrected |

\* Hashimoto & Sakai, 2003 had a total sample size of 18, 15 of which participated in the experiment used in the meta-analysis. Participant demographics were only given for the full sample size ( $n = 18$ ). Details pertaining to how they corrected for multiple comparisons were not given.

**Table S2.**
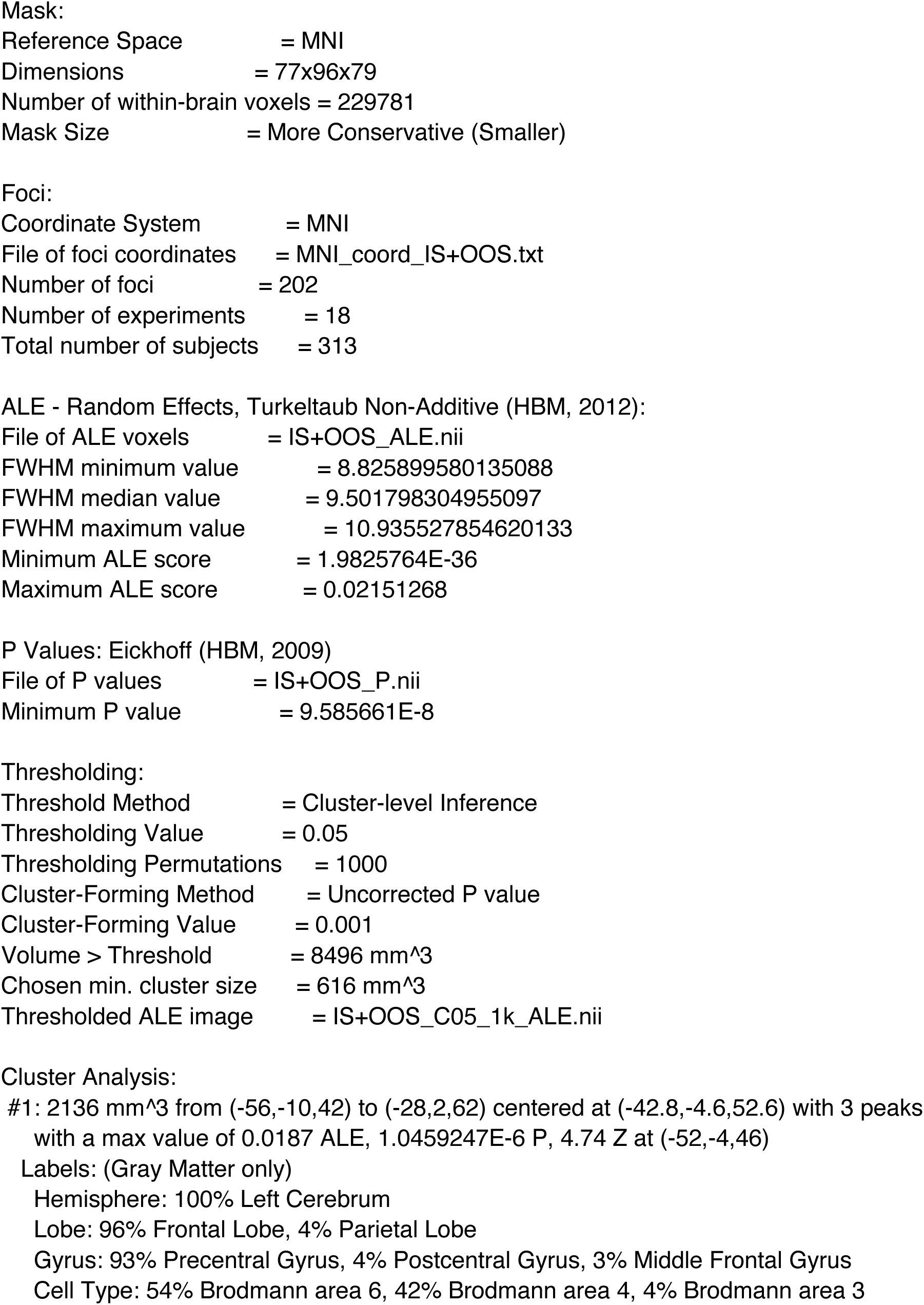
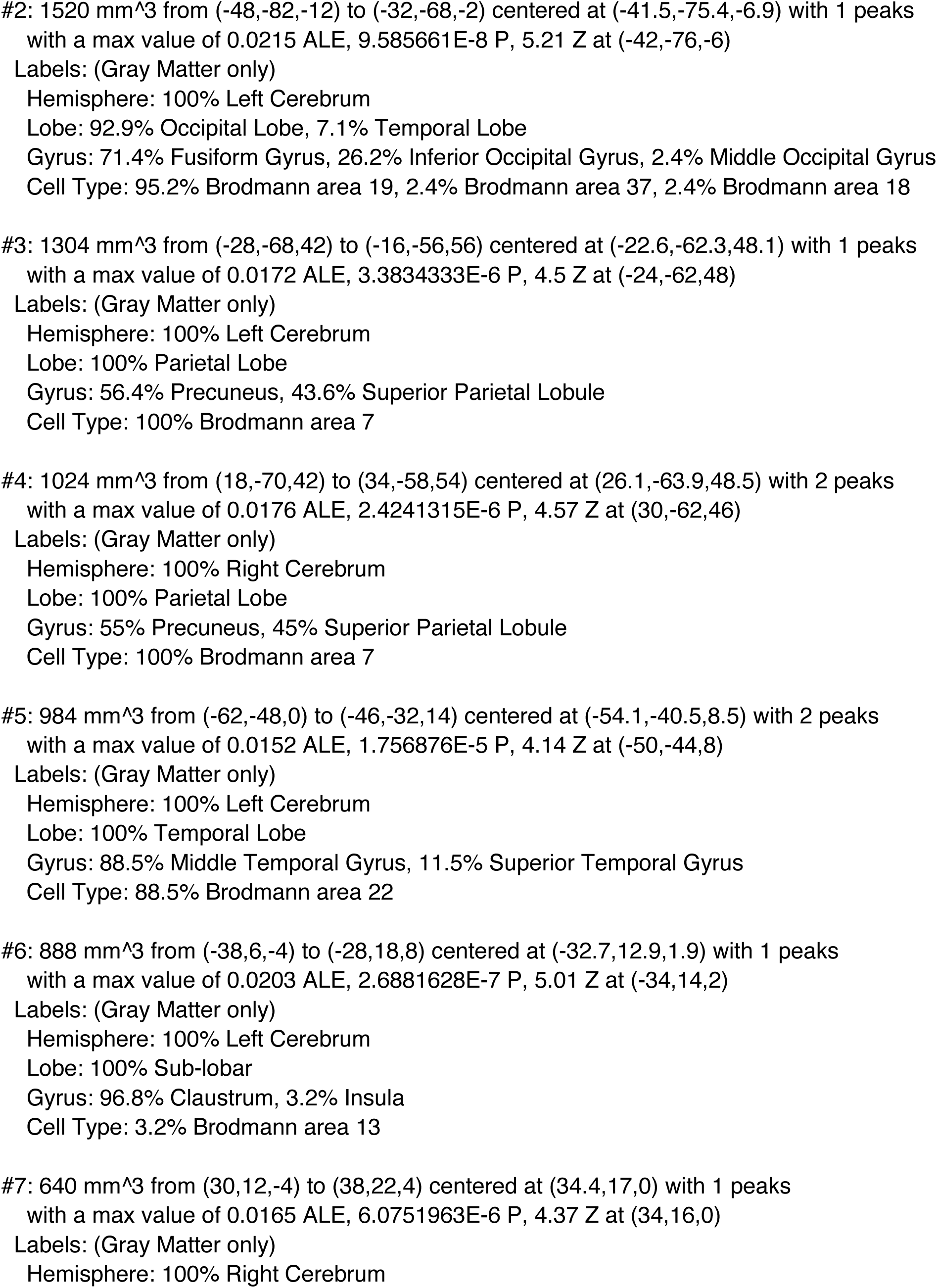
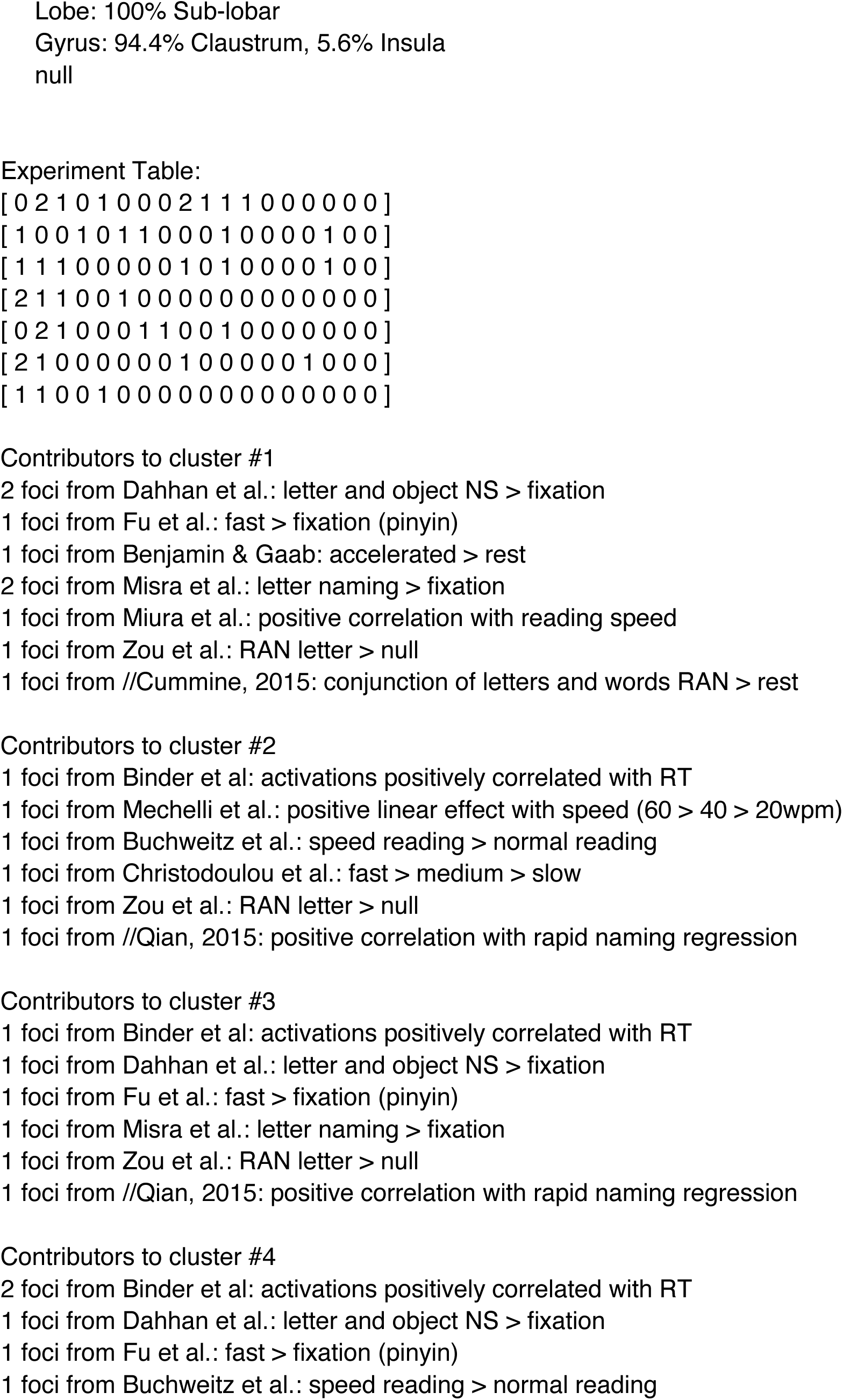
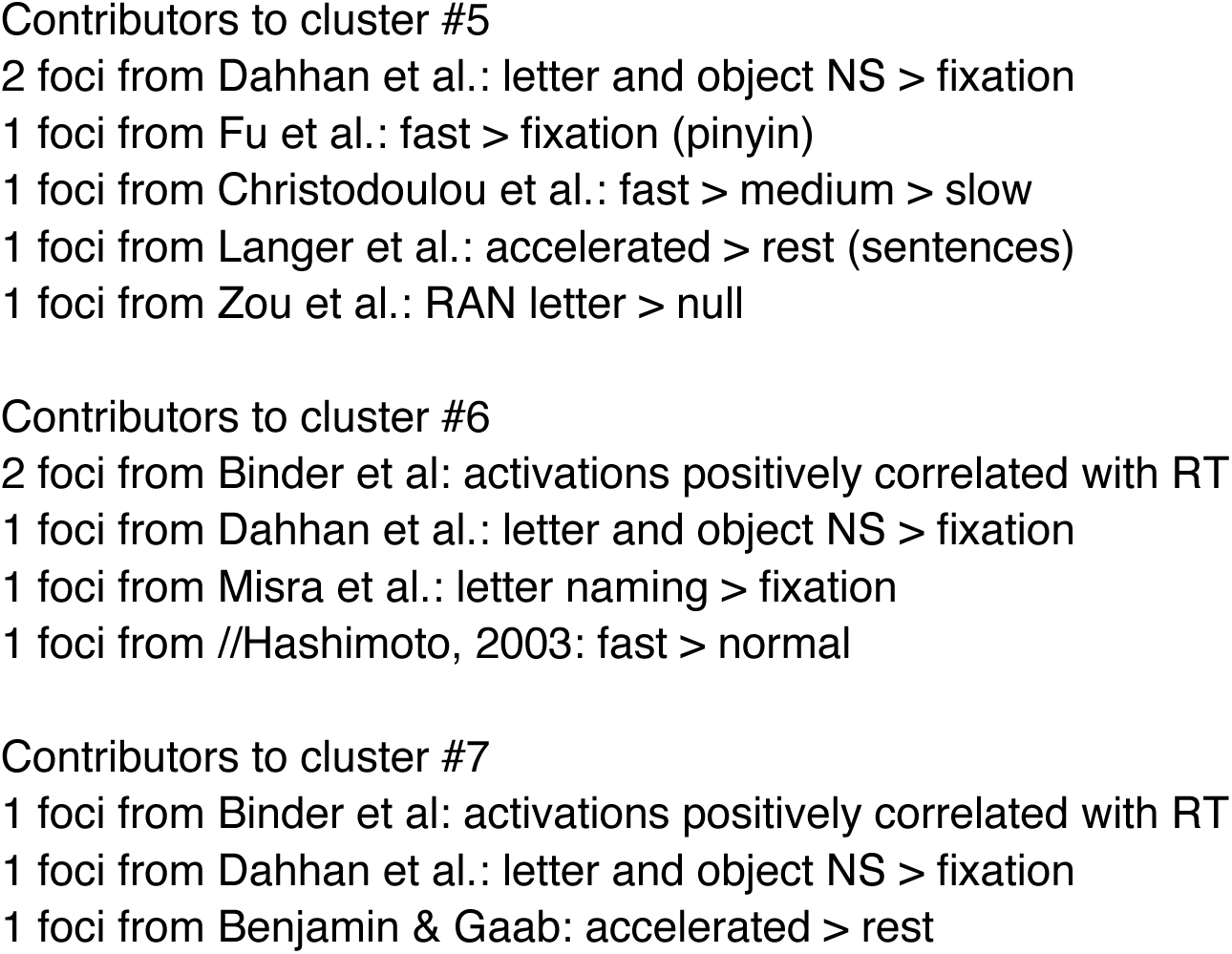
GingerALE output.

